# CAD variants act through ox-LDL-induced enhancer remodelling to alter VSMC gene programmes

**DOI:** 10.64898/2026.04.21.718843

**Authors:** Thomas A. Agbaedeng, Goutham Atla, Thomas K. Hiron, Jiahao Jiang, Yashaswat Malhotra, Lewis Marsh, Joanna M.M. Howson, Christopher A. O’Callaghan

**Affiliations:** Nuffield Department of Medicine, University of Oxford, Oxford, UK; Novo Nordisk Research Centre Oxford, UK

## Abstract

**Background:** Vascular smooth muscle cells (VSMCs) play a central role in atherosclerotic coronary artery disease (CAD). Oxidised low-density lipoprotein cholesterol (ox-LDL) induces VSMCs dysfunction but the underlying molecular mechanisms are unclear. CAD genome-wide association studies (GWAS) have identified hundreds of disease-associated loci but their biological roles remain poorly defined. We hypothesised that ox-LDL drives pro-atherogenic changes in VSMCs by altering gene regulatory programs involving causal CAD variants.

**Methods:** *Ex-vivo* human coronary VSMCs were exposed to ox-LDL and profiled using RNA-seq, ATAC-seq, and H3K27ac ChIPmentation. Enhancer-gene links were inferred by integrating these data with Hi-C using the Activity-by-Contact (ABC) model. Variant effect predictions were done using AlphaGenome and key target genes functionally tested by CRISPR/Cas9 knockout.

**Results:** Ox-LDL induced widespread transcriptional reprogramming in coronary VSMCs, with 1,487 upregulated and 1,864 downregulated genes (FDR < 0.05). Single-cell RNA-seq meta-analysis demonstrated that ox-LDL-associated programmes enriched in pro-inflammatory and synthetic-inflammatory VSMC clusters *in vivo*. ATAC-seq identified ∼22k differentially accessible regions following ox-LDL exposure (FDR < 0.05). Integration of ATAC-seq, H3K27ac, and Hi-C using the ABC framework showed that ox-LDL-driven chromatin remodelling was concentrated at distal enhancers, which linked to 2,008 differentially expressed genes via 4,243 peak–gene connections. ABC enhancers were significantly enriched for CAD variants compared with non-vascular disease controls, with stronger enrichment in dynamically accessible enhancers. AlphaGenome predicted larger regulatory effects of prioritised CAD variants in smooth muscle cells than in a non-vascular comparator, and motif analyses indicated allele-dependent transcription factor binding at prioritised enhancer variants. Locus-level prioritisation nominated candidate enhancer-mediated mechanisms at the *SPECC1L* and *MAP1S* loci, and CRISPR knockout of the target genes *GUCD1* and *BACH1* rescued ox-LDL-induced growth arrest/senescence phenotypes in human coronary artery VSMCs.

**Conclusions:** Our unbiased multi-omics framework shows that ox-LDL rewires VSMC regulatory programmes that influence CAD genetic risk. Enhancer–gene mapping refines effector-gene assignment at CAD loci and prioritises regulatory targets in coronary VSMCs.

## INTRODUCTION

Atherosclerotic cardiovascular disease, such as coronary artery disease (CAD), stroke, and peripheral artery disease, remain the leading causes of disability and death worldwide.^1^ This is a significant burden for individuals, healthcare systems, and economies. A rising incidence of CAD has been attributed to the growing prevalence of cardiometabolic risk factors, including elevated low-density lipoprotein (LDL) cholesterol levels.^1,2^

The pathophysiology of atherosclerosis is a multifaceted process involving the accumulation of lipids and cells in plaques which narrow the vessel lumen.^3^ Recent genetic fate-mapping studies have revealed extensive vascular smooth muscle cells (VSMCs) plasticity in plaques and its role in lesion formation. Within atherosclerotic lesions, oxidised LDL-cholesterol (ox-LDL) is taken up by macrophage resulting in a ‘foamy’ phenotype.^4^ Ox-LDL has other effects on VSMCs including loss and disorganisation of cytoskeletal proteins^5^, altered extracellular matrix synthesis^6^, and acquisition of inflammatory phenotypes ^7^. Both *in vitro* experiments and analysis of human plaques indicate that VSMCs can express the scavenger receptor CD68, a biomarker widely used to identify macrophages.^8,9^ However, the mechanisms driving ox-LDL-induced VSMC dysfunction are not clear.

Genome-wide association studies (GWAS) have identified over 393 genetic variants associated with CAD.^10,11^ Subsequent polygenic prediction modelling indicates that CAD risk results from contributions by many, low impact genetic variants.^12^ Enrichment of CAD GWAS SNPs in VSMCs has been reported.^13^ However, the mechanisms by which these variants influence disease risk remains undefined because most disease-associated variants occur in non-coding regions, particularly in potential regulatory elements such as enhancers or promoters.^14 11^ Integrating epigenomic data with GWAS results has led to the identification of functional roles for likely causal variants operating in the macrophage and endothelial cell response to ox-LDL.^15,16^

The response of VSMC to ox-LDL and the role of this interaction in human CAD is incompletely understood. We hypothesised that ox-LDL drives pro-atherogenic changes in VSMCs by altering gene regulatory programs involving causal CAD variants. To test this hypothesis, we generated genome-wide maps of the chromatin accessibility, regulatory activity, and gene expression changes induced by ox-LDL in primary human coronary artery VSMC using RNA-seq, ATAC-seq, and ChIPmentation. We integrated these genome-wide chromatin maps with genetic information from CAD GWAS to prioritise causal CAD SNPs in VSMCs.

## METHODS

### Cell Culture

Primary human coronary artery smooth muscle cells (Human coronary artery VSMCs) were obtained at passage 2 from Promocell (Heidelberg, Germany) and expanded before usage. The catalogue numbers for the six HCASMC lines used in this study are: #440Z021.2, #451Z031.6, #452Z013.3, #458Z016.8, #482Z028.2, and #488Z024.3. All donors were used at passage 4 for experiments. Human coronary artery VSMCs were maintained in Smooth Muscle Cell Growth Medium MV2 (Promocell) supplemented with 100 units/mL penicillin and 100 µg/mL streptomycin (Sigma-Aldrich, St. Louis, MO, USA). Cells were assayed at ≥90% confluency at passage 4.

### Cell Treatments

LDL was purified from human plasma by isopycnic ultracentrifugation, then oxidized overnight using 25 µM CuCl_2_ for 18 h, as described previously. DPBS solution (Thermo Scientific, Waltham, MA, USA) from the last round of dialysis was used as the control buffer. The concentration of ox-LDL was determined using the BCA protein assay (Thermo Scientific). Unless otherwise specified, cells were treated with a final concentration of 100 ug/mL of ox-LDL or an equal volume of control buffer for 48 h.

### Library Preparation for Bulk RNA-seq

Total RNA from Human coronary artery VSMCs was purified using RNeasy Micro Kit (Qiagen, Hilden, Germany) after TRIzol and chloroform extraction according to the manufacturer’s protocol. The quality of RNA samples was analysed using a TapeStation 2200 with High Sensitivity RNA ScreenTape (Agilent Technologies, Santa Clara, US) and nanodrop. Samples with high integrity (RIN score>9) were selected for PolyA-enrichment and TruSeq library preparation (Illumina, Inc., San Diego, CA, USA). Samples were sequenced to a target depth of 30 million reads as 75-bp paired-end reads using a NovaSeq 6000 device (Illumina).

### Library Preparation for ATAC-seq

Libraries for ATAC-seq were prepared according to the Omni-ATAC protocol.^17^ In brief, ∼50,000 viable cells were pelleted and incubated in ATAC-Resuspension Buffer (ATAC-RSB) containing 0.1% NP40 (Roche, Basel, Switzerland), 0.1% Tween-20 (Roche, Basel, 584 Switzerland) and 0.01% Digitonin (Promega, Madison, WI) on ice for 3 min to isolate nuclei. After washing in 1 mL of ATAC-RSB containing 0.1% Tween-20, nuclei were tagmented by Tn5 at 37 °C for 30 min in a thermomixer (1000 rpm, Eppendorf Thermomixer F1.5, Sigma-Aldrich) using the Nextera kit (Illumina), then they were cleaned up using Zymo DNA Clean and Concentrator-5 Kit (Zymo Research, Orange, CA, USA). Samples were then PCR amplified using NEBNext High-Fidelity PCR master mix (New England Biolabs, Ipswich, MA, USA) and index primers from Nextera Kit (Illumina), and analysed using a TapeStation 2200 with High Sensitivity D1000 ScreenTape (Agilent). Libraries showing successful transposition were sequenced to a target depth of ≥100 million read depth, 75-bp read pairs using a NovaSeq X instrument (Illumina).

### Library Preparation for ChIPmentation

Libraries for ChIPmentation were performed as previously described by Schmidl et al^18^ with modifications. Cells were harvested, counted, and resuspended in Complete Medium. Cells (∼100k) were fixed by adding fresh paraformaldehyde to a concentration of 1% for 10 min with mixing, then quenched by adding glycine to a final concentration of 0.125 M for 5 min all at 20–25 °C. After washing in ice-cold PBS containing 1 µM phenylmethylsulfonyl fluoride (PMSF, Abcam, Cambridge, UK), cells were sonicated in a buffer containing 10 mM tris-HCl (Thermo Fisher), 0.25% SDS (sodium dodecyl sulfate, Thermo Fisher), 2 mM EDTA (Insight Biotechnology Ltd, Wembley, UK), 1× cOmplete^TM^ protease inhibitor cocktail (PIC, Sigma-Aldrich), 1 µM PMSF, and sterile H_2_O using 16 cycles of 30 s ON/30 s OFF. Following shearing, a quarter of the lysate was immunoprecipitated using 1 µg of H3K27ac antibody (Abcam) and 1% was preserved as an input control. Immunoprecipitated chromatin samples (IP) and input controls were incubated and tagmented in a tagmentation buffer containing 10 tris-HCl, 5 mM MgCl_2_ (Thermo Fisher), 10% N,N-dimethylformamide (Sigma-Aldrich), 1 µL Tn5 from the Nextera kit (Illumina), and sterile H_2_O for 10 min in a thermomixer (800 rpm, Sigma-Aldrich), then purified using Qiagen MinElute kit. Tagmentation reactions were quenched with two washes in RIPA lysis (RIPA-LS) buffer containing 50 mM tris-HCl, 2 mM EDTA, 0.1% SDS, 0.1% sodium deoxycholate (NaDOC), 1% Triton x-100 (Sigma-Aldrich), and sterile H_2_O followed by a single wash in buffer containing 10 mM tris-HCl, 1 mM EDTA, and sterile H_2_O. IP samples and input controls were de-crosslinked in ChIP Elution Buffer containing 10 mM tris-HCl, 5 mM EDTA, 300 mM NaCl, 0.4% SDS, 1 µL Proteinase K (ThermoFisher), and sterile H_2_O by incubating at 55 °C for 1 h and 65 °C for ∼6 h in a thermomixer (800 rpm), followed by treatment with 1:100 RNase A solution (Sigma-Aldrich) at 37 °C for 30 min, then cleaned up with MinElute (Qiagen). Chromatin samples were then PCR amplified using NEBNext High-Fidelity PCR master mix (New England Biolabs) and index primers from Nextera Kit (Illumina), and analysed using a TapeStation 2200 with High Sensitivity D1000 ScreenTape (Agilent). Libraries were filtered to retain those with fragment sizes peaking at 200 bp by SPRI right-sided size selection (Beckman Coulter, Brea, CA, USA), then they were sequenced to a target depth of ≥30 million read depth, 75-bp read pairs using a NovaSeq X instrument (Illumina).

### Data Analysis for RNA-seq

Sequenced reads were first quality checked using fastQC v0.11.9 and compiled using MultiQC^19^, then trimmed to remove sequencing adaptors using NGmerge v0.3^20^. Trimmed reads were aligned to the reference human genome (GRCh38.p14) using the splice-aware STAR aligner (v2.7.10b).^21^ Human genome assembly GRCh38.p14 and gene annotation release 44 were downloaded from the GENCODE site and were indexed using STAR. Unmapped or multi-mapping reads were filtered out to retain uniquely mapped reads using SAMtools v0.1.19.^22^ Gene-wise count matrices were generated using the featureCounts function of the Rsubread package (v2.12.3).^23^ Differential expression analysis was performed in R using the edgeR package.^24^ Genes with expression of <50 read counts in more than half of the samples were filtered out, then gene count matrices were normalised by sequencing depth and by the Trimmed Mean of M-values (TMM) method. The quasi-likelihood test was used to perform differential gene expression between ox-LDL versus buffer groups, with biological donor effects controlled using a generalised linear model, with the Benjamini-Hochberg (B-H) method used to adjust for multiple comparisons. Differentially expressed genes (DEGs) were identified as genes with adjusted p-values *(P.adj*) <0.05. DEGs were subjected to pathway enrichment analyses, including gene ontology (GO) and Kyoto Encyclopaedia of Gene and Genome (KEGG), using the clusterProfiler package.^25^

### Data Analysis for ATAC-seq

Sequenced ATAC-seq reads were first quality checked using fastQC and compiled using MultiQC^19^, then they were trimmed to remove adaptor reads using TrimGalore v0.6.10 (URL: https://github.com/FelixKrueger/TrimGalore/blob/master/Docs/Trim_Galore_User_Guid <u>e.md</u>). To align reads to human genome assembly hg38, a Bowtie index was built using Bowtie2^26^ with the genome build downloaded from USFC Genome Browser. Trimmed reads were aligned to the hg38 reference genome using Bowtie2 with *--very-sensitive* settings, sorted by coordinates and indexed with SAMtools. PCR duplicates were marked using Picard^27^ and, together with mitochondrial reads, were removed from downstream analysis using SAMtools. Post-alignment quality control (QC) was performed as instructed by the ENCODE ATAC-seq processing pipeline, retaining libraries with Non-Redundant Fraction (NRF) >0.9, PCR Bottleneck Coefficients (PBC)-1 >0.9, PBC2 >3, and Transcription Start Site (TSS) Enrichment score >7. For downstream analysis, reads were shifted using BEDTools v2.30.0 to account for the Tn5-introduced DNA nicks, then a consensus peak set was called using MACS3 combining all sequenced samples per treatment group and for all groups. Peaks overlapping the ENCODE blacklisted region were removed using BEDTools before counting mapped reads with RSubread. Differential accessibility analysis was performed using edgeR as described above, with both within- and between-lane normalisations performed to offset GC bias using EDAseq and qsmoothGC.^28^ Differentially accessible peaks (DAPs) were identified using *P.adj* <0.05, then annotated using ChIPseeker^29^ followed by pathway enrichment analyses as previously described.

### Data Analysis for ChIPmentation

Sequenced reads were first quality checked using fastQC and compiled using MultiQC^19^, then they were trimmed to remove adaptor reads using TrimGalore. Trimmed reads were aligned to the hg38 reference genome using Bowtie2 with *--very-sensitive* settings, sorted by coordinates and indexed with SAMtools. PCR duplicates were marked using Picard^27^ and removed from downstream analysis using SAMtools. Post-alignment QC was performed as instructed by the ENCODE ChIP-seq processing pipeline, retaining libraries with NRF >0.9, PBC1 >0.9, and PBC2 >10. For downstream analysis, reads were shifted using BEDTools v2.30.0 to account for the Tn5-introduced DNA nicks, then a consensus peak set was called using MACS3 combining all sequenced samples per treatment group and for all groups. Peaks overlapping the ENCODE blacklisted region were removed using BEDTools before counting mapped reads with RSubread. Differential accessibility analysis was performed using edgeR as described above, with both within- and between-lane normalisations performed to offset GC bias using EDAseq and qsmoothGC.^28^ Peaks with differential H3K27ac activities/signals were identified using *P*.adj <0.05, then annotated using ChIPseeker^29^ followed by pathway enrichment analyses as previously described.

### Single-cell RNA-seq (scRNA-seq) analysis of human coronary plaque VSMCs

We retrieved sequencing data of 7 datasets of scRNA-seq datasets on human coronary plaques and non-plaque control artery tissues reported in two studies from the Gene Expression Omnibus (GEO) using accession number GSE131780^30^ and from Zenodo using 6032099^31^. Sequencing reads were re-aligned against the GRCh38 reference genome using Cell Ranger (v5.0; 10x Genomics, Pleasanton, CA). Count matrices were then aggregated in Seurat V5^32^. The following thresholds were used for removing low-quality cells: detected genes <200, read counts < 500, percent mitochondrial reads (mt%) > 20%, and doublet rates of 5%. Samples were then integrated using Harmony and subjected clustering analysis (n_PC = 18, k.param = 20, and resolution = 0.1). We next performed cell annotations using singleR package^33^ and selected the VSMC object for further analysis. For subclustering of VSMCs, the number of principle components (PCs), the k value for the k-nearest neighbour (KNN) graph, and clustering resolution were optimized using the scclusteval package^34^. Differential gene expression analysis was performed using egdeR, using TMM-normalised counts.

### Hi-C data analysis

Raw Hi-C data from Human coronary artery VSMCs were retrieved from GEO using the accession number GSE141749^35^. Paired-end sequencing reads were processed and aligned using HiCUP (v0.9.2)^36^ pipeline. Briefly, paired-end reads were aligned to the GRCh38 reference genome within the HiCUP pipeline, producing filtered, coordinate-sorted alignment files. The resulting BAM file was then streamed with SAMtools and converted to a de-duplicated paired-contacts file using pairtools (v1.1.3)^37^. For Hi-C matrix generation, the de-duplicated pairs were reformatted to the Juicer short input format and processed with Juicer Tools (v1.6)^38^ to build a multi-resolution contact map using Knight–Ruiz (KR) normalisation.

### Enhancer-target gene prediction

We used the Activity-by-contact (ABC) model to construct regulatory map of enhancer-promoter interactions to link active enhancers to their target gene, as previously described.^39^ Briefly, we built enhancer-gene maps by integrating chromatin accessibility (ATAC-seq), H3K27ac ChIPmentation, and chromatin conformation capture (Hi-C) contact frequencies from human coronary artery VSMCs.^35,39,40^ First, MACS2 was used to call peaks in accessible chromatin with a lenient *P*-value <0.1, then reads were counted, retaining the top 150,000 peaks with the highest read counts. Next, peaks were resized to 500 bp centred on their summits, a further set of peaks of 500 bp centred on all gene TSS’s added, and those overlapping blacklisted regions excluded. A final set of candidate regions was created by merging any overlapping peaks. The element activity was calculated by counting the reads in each candidate region from both ATAC-seq and H3K27ac ChIPmentation datasets, then the geometric mean of the two assays was taken. Fourth, the ABC score was computed as product of element activity × KR-normalised Hi-C contact, then normalised this score product by sum of products of activity × contact for all other nearby elements in ±5 Mb region the target gene. The best ABC score threshold for identifying significant enhancer-gene regulatory interactions was automatically computed. Enhancer-gene pairs with ABC score of ≥0.015 were used for this analysis.

### Multi-omics Integration Analyses

*Cis*-regulatory elements (CREs) were defined by intersecting quantile normalised ATAC-seq peaks with candidate regions from ABC enhancer-gene maps using the *findOverlaps* function of the GenomicRanges package v1.56.2^41^. CRE peaks were annotated using ChIPseeker, with promoters defined as regions within ±500 pb of TSS and enhancers defined as active regions not annotated to promoters, including intronic and distal elements. Dynamic CRE peaks as those with ATAC-seq *P*.adj <0.05, then subjected to pathway enrichment analyses as previously described. The concordance between chromatin accessibility and H3K27ac signal at dynamic CREs was assessed using Pearson’s product moment correlation and plotted using ggplot2. Regulatory activity at ABC defined CREs was computed as: the sum of CRE ATAC-seq log_2_FC, -log_10_ of CRE ATAC-seq FDR, RNA-seq log_2_FC, and - log_10_ RNA-seq FDR. Differential accessibility enhancers of ABC-linked DEGs versus non-DEGs were tested using the two-sample Wilcoxon test.

### Transcription factor (TF) motif enrichment and footprinting analysis

Transcription factor motif enrichment analysis was performed using default settings of the Analysis of Motif Enrichment (AME) tool^42^ of the MEME-Suite^43^ and the JASPAR2024^44^ database. Peaks were first converted from BED to FASTA file formats using BEDtools. Motif enrichment across peaks was performed using a set of background, non-differential peaks as controls for enhancers or non-promoter peaks, or shuffled promoter background peaks of the same length for promoters. TF motifs were prioritised based on expression in the RNA-seq dataset, direction of expression, and significance. Thus, ox-LDL TFs were upregulated DEGs enriched in ox-LDL upregulated dynamic peaks, whereas buffer TFs were downregulated DEGs enriched in ox-LDL repressed peaks.

Footprint analysis or scanning for occurrences of prioritised TF motifs in peaks were performed using the FIMO (Find Instances of Motif Occurrence) too^l45^ of MEME-Suite. Genome-wide TF footprinting on raw consensus ATAC-seq reads per condition was performed using the TOBIAS (Transcription factor Occupancy prediction By Investigation of ATAC-seq Signal) suite.^46^

### GWAS Enrichment Analysis

GWAS statistics were downloaded from the NHGRI-EBI Catalog database (October 2024). SNPs with significant genome-wide association (*P* <5 × 10^-8^) were pruned (r^2^ > 0.1 identified by 1000 Genome Project phase 3 in European [EUR] samples, hg38 build) to remove redundant loci, and then all SNPs in high linkage disequilibrium (LD) within each locus (r^2^ > 0.8 in EUR samples) were used to establish a set of trait-associated variants using PLINK v1.90b6.26 70. Only dbSNP common variants (minor allele frequency [MAF] >1%, dbSNP database version 155) were considered in this study.

To assess overlap of GWAS SNPs with regulatory regions, we identified SNPs intersecting ABC-linked CRE peaks in human coronary artery VSMCs using the *findOverlaps* function. We then compared overlap of CAD-associated SNPs with overlap of SNPs associated non-vascular inflammatory conditions, including inflammatory bowel disease (IBD), systemic lupus erythematosus (SLE), and rheumatoid arthritis (RA), using Fisher’s exact test. We also compared overlap of CAD SNPs in ABC-linked peaks with overlap in ATAC-seq peaks that were not linked by ABC to any expressed genes in VSMCs.

To assess the significance of CAD SNP enrichment in chromatin accessibility peaks and ABC-linked regulatory regions, we used GREGOR (Genomic Regulatory Elements and Gwas Overlap algoRithm).^47^ For all the GWAS lead SNPs, GREGOR selects matched control SNPs from across the genome that match for (i) number of variants in LD, (ii) minor allele frequency, and (iii) distance to the nearest gene. The observed number of index SNPs (or their LD proxies, r² ≥ 0.8, EUR) overlapping a given set of genomic regions is then compared to the expected overlap derived from the matched control sets, with significance assessed using a binomial-based test. ATAC-seq consensus peaks, differential ATAC-seq peaks, ABC-linked peaks, and differential ABC-linked peaks were tested for enrichment of CAD GWAS index SNPs. To confirm that enrichment was specific to biologically meaningful regulatory regions, size-matched random genomic regions were also evaluated as negative controls.

### GWAS Credible Set and Fine Mapping Analysis

Credible set fine-mapping was performed using CAD GWAS summary statistics from Aragam et al. Briefly, variants reaching genome-wide significance (P < 5 × 10^-8^) were selected as seed signals and grouped into loci by aggregating all significant variants within ±5 Mb of each seed; overlapping windows were then merged iteratively until all loci were non-overlapping. For each locus, we computed LD using a reference panel of 10,000 unrelated UK Biobank participants of EUR ancestry. LD was calculated in PLINK v1.9 as the Pearson correlation (r) between variants, and analyses were restricted to variants present in the LD panel. We applied SuSiE-RSS using locus-specific z-scores and the pre-computed LD matrix to derive 95% credible sets. Fine-mapping was run with the susieR package^48^, using coverage = 0.95 and a purity threshold of 0.5 (min_abs_corr = 0.5), with all other parameters left at default settings.

### Genotyping and variant calling

Aligned ATAC-seq BAM files were used for variant discovery, and variants were called per sample using GATK HaplotypeCaller in GVCF mode.^49^ Per-sample GVCFs were merged using CombineGVCFs, and joint genotyping was performed with GenotypeGVCFs to generate a multi-sample VCF. Variant files were indexed with tabix and validated using ValidateVariants (with --validate-GVCF for GVCFs). The resulting joint-called VCF was used as input for downstream allele-specific analyses.

### *In Silico* Prediction of the Effects of CAD SNPs

Genome-wide regulatory effects of non-coding variants were predicted using AlphaGenome, a deep-learning model developed by DeepMind that jointly models sequence-based regulation across diverse molecular assays and cell types. Breifly, a 1 Mb genomic window centred on each variant was provided as input to the model. AlphaGenome uses a convolutional–transformer architecture, in which convolutional layers learn local sequence features such as transcription factor motifs and splice signals, followed by transformer blocks that capture long-range regulatory interactions across hundreds of kilobases. The shared backbone feeds into multi-task output heads trained on large-scale functional genomics datasets (including ENCODE, GTEx, and FANTOM), enabling simultaneous prediction of chromatin accessibility (ATAC-seq/DNase-seq), histone modifications, TF binding, transcriptional output (RNA-seq, CAGE/PRO-cap), splicing, and 3D chromatin contact maps.

Variant effects were computed by comparing predictions for reference and alternate alleles at each locus, using task-specific aggregation functions. Predicted scores were summarized per assay and biospecimen and used for downstream prioritization, comparison across cell types, and integration with enhancer–gene links derived from ABC modelling.

We determined disruptive effects of non-coding variants on TF motifs by performing motif breaking analysis using the MotifbreakR package^50^. Briefly, our list of prioritised SNPs were read from dbSNP155 to obtain locations using SNPlocs and reference sequence was determined using BSgenome, all in human genome build hg38. We queried position-weighted matrices (PWM) of TF binding motifs from JASPAR Core and HOCOMOCOv11 Core A–C via MotifDb. We employed the relative entropy method to calculate frequency scores for the reference and alternative alleles. SNPs with either allele scores above the threshold 0.85 were reported and filtered to retain only significant PWM matches (*P* <0.0001). We further filtered SNPs to retain those strongly affected TFBSs and calculated the alternative allele’s binding ability difference (ΔPWM). Motifs with -ΔPWM were considered disrupted and those with +ΔPWM were considered to be created.

### Genome-wide Transcription Factor Activity

ATAC-seq data from Human coronary artery VSMCs treated with ox-LDL or buffer control were analysed using the chromVAR package to quantify global TF activity. Motif PWMs were obtained from the JASPAR2022 database. Differential motif accessibility was calculated as the change in deviation scores between ox-LDL vs Buffer (deviation), with positive values indicating increased global TF occupancy under stress.

### Allele-specific Accessibility Analysis

For illustrative candidate loci, allele-specific accessibility was explored using ATAC-seq read counts at heterozygous SNP positions identified from the jointly called VCF. Reference and alternate allele depths were extracted from the AD field in the VCF, and mean allele-specific depth was summarised within each condition after applying a minimum depth threshold of 10 reads per site. Because heterozygosity at some prioritised loci was observed in only a single donor, these analyses were considered descriptive and were not used as primary evidence for candidate prioritisation.

### Variant-to-Motif-to-Target Gene Mapping

To identify specific TF-SNP-target gene relationships, variants were first intersected with ABC-linked regulatory elements to assign predicted target genes. Candidate loci were then scanned for motif disruption using the motifmatchr and the JASPAR PWM library within a 50-bp window centred on each SNP. Motif matches were integrated with chromVAR differential deviation scores, RNA-seq expression of the corresponding TFs, and target-gene differential expression. TFs were prioritised if they were expressed in human coronary artery VSMCs and had significant global activity changes (chromVAR, deviation).

### CRISPR/Cas9 mediated gene knockout

For CRISPR/Cas9-mediated gene knockouts, guide RNAs (gRNAs) were designed using the Invitrogen TrueDesign Genome Editor (https://www.fisher.com/uk/en/home/life-science/genome-editing/invitrogen-truedesign-genome-editor.html). All oligonucleotides were ordered via IDT, then single-guide RNAs were synthesised via *in-vitro* transcription (IVT) using the Precision gRNA Synthesis Kit (Thermo Fisher) according to manufacturer’s recommendation. All oligonucleotides used are shown in **Table S1**. For transfection, Human coronary artery VSMCs were seeded at the density of 1.5 × 10^5^/well in 6-well plates for 24 h. Transfection was performed in Human coronary artery VSMCs at ∼70% confluency by co-transfecting 4.0 μg of TrueCut Cas9 protein (v2, Thermo Fisher) and 1.0 µg of IVT sgRNA using Lipofectamine™ CRISPRMAX™ Cas9 Transfection Reagent (Thermo Fisher) for 48 h.

### Flow cytometry

Treated cells were detached using StemPro Accutase solution (Thermo Scientific) and re-suspended in ice-cold PBS. Cells were stained with Zombie Violet (423113, Biolegend) at a final dilution of 1:200 for cell viability followed by washes in ice-cold PBS. Cells were fixed in 2% PFA and permeabilised in ice-cold FACS PERM buffer (PBS, 1% BSA, 0.1% Saponin, and 1% sodium azide). Primary antibody staining was carried out at 4⁰C for 30 min with 1:150 Mouse anti-human p21 (353102, Biolegend). Secondary antibody staining was performed with Rat anti-mouse FITC at a dilution of 1:500 (406001, Invitrogen) at 4⁰C for 30 min. Data was acquired on FACSCanto II Flow Cytometer (BD Biosciences, East Rutherford, NJ, USA) and exported to FlowJo 10 (FlowJo LLC, Ashland, OR, USA) for quantitative analysis.

### Cell Proliferation

For cell proliferation, cells were washed twice in PBS and stained in CellTrace™ (CellTrace™ CFSE Cell Proliferation Kit, ThermoFisher) for 20 min at 20–25 °C following manufacturer’s recommendation. Stained cells were seeded and treated with ox-LDL or Buffer for 4 days to track cell divisions. At the end of the experiments, cells were harvested as previously described and prepared for flow cytometry.

### Data Visualisation

To visualise ATAC-seq, ChIPmentation, and RNA-seq data, aligned reads in BAM formats were normalised to counts per million with a binSize of 100 using deepTools v3.5.5^51^ and converted to bigwig files. Bigwig files, consensus peaks in BED format, and enriched SNPs in BED format were visualised in Integrated Genome Browser v2.16.0.^52^ All plots were generated using ggplot2 or functions adapted from ggplot2 in R v4.4.1 (R Foundation for Statistical Computing, Vienna, Austria).

## RESULTS

### Ox-LDL drives widespread transcriptomic reprogramming in coronary VSMCs

We first assessed ox-LDL uptake and found that ox-LDL uptake was evident in Human coronary artery VSMCs within 48 h (**Fig-1B**). We performed RNA-seq to analyse the transcriptomic effects of ox-LDL on Human coronary artery VSMCs (**Fig-1A**). Exposure to ox-LDL led to a significant change in the expression of 3,351 of the 12,409 genes expressed in Human coronary artery VSMCs (*P*.adj <0.05, **Fig-1C**), with 1,487 genes upregulated and 1,864 downregulated. Ox-LDL upregulated several lipid-handling and proinflammatory genes, including *AKR1B10*, *LACC1*, and *SPP1*, but downregulated atheroprotective genes such as *KLF2* and *MGP,* and proliferative genes such as *KIF20A* and *RASL11A* (**Fig-1D**). KEGG and GO enrichment analyses demonstrated that genes upregulated by ox-LDL are enriched for lipid-related pathways in atherosclerosis, autophagy, HIF-1 signalling and ferroptosis pathways (*P*.adj <0.05, **Fig-1E**, **Fig-S2**, **Table S2**). In contrast, downregulated genes showed significant enrichment of extracellular matrix, cell division and proliferation, negative regulation of cell migration, and cellular senescence pathways (*P*.adj <0.05, **Fig-1E**, **Fig-S2**, **Tables S2–3**).

**Fig-1.**
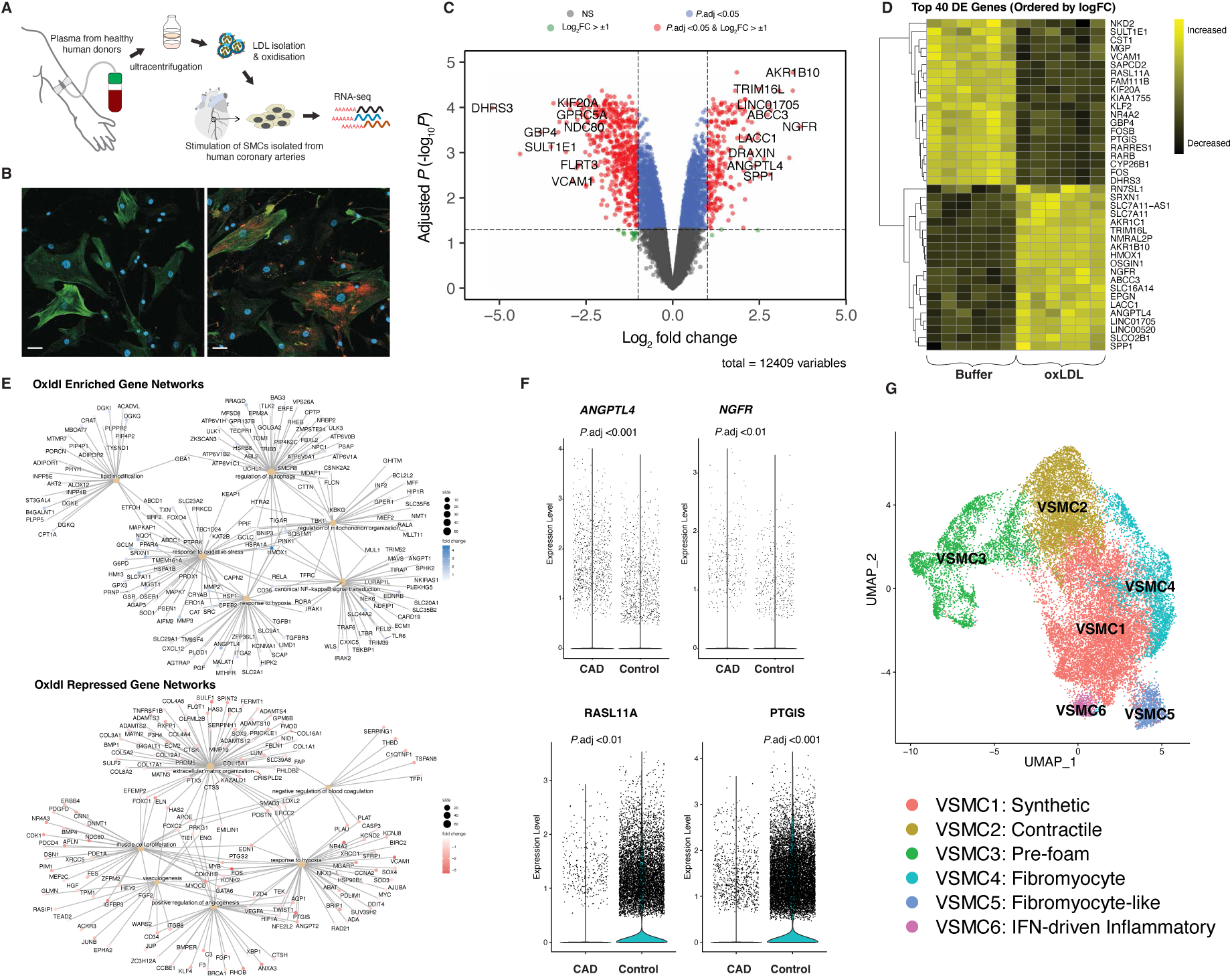
Transcriptomic changes induced by ox-LDL in human coronary artery VSMCs overlap with those in coronary plaque VSMC. **A**, Experimental workflow for transcriptomic analysis. **B**, Representative super-resolution confocal microscopy images of coronary artery VSMCs exposed to buffer (left) or DiI-labelled ox-LDL (right). Scale bar: 200 µm; Red, DiI-ox-LDL; Green, α-SMA; Magenta, DAPI. **C**, Volcano plot of the transcriptomic changes induced by ox-LDL compared to buffer. The horizontal dashed line represents P.adj <0.05 and vertical dashed lines are log_2_ fold change (log_2_FC) = -1.0 or +1.0. NS, not significant; P.adj, p-values adjusted for multiple comparison using the Benjamini-Hochberg false-discovery rate (FDR) method. **D**, Heatmap of the transcriptomic profiles and top 40 differentially expressed (DE) genes influenced by ox-LDL compared to buffer. **E**, Network maps of interactions between gene ontology (GO) terms of genes with higher expression (upper panel) or lower expression (lower panel) when exposed to ox-LDL compared to buffer. **F**, Expression of some of the top DEGs induced by exposure to ox-LDL perturbation experiments in single-cell RNA-sequence (scRNA-seq) of coronary plaques of people with CAD vs normal arteries from people without CAD. **G**, Uniform manifold approximation and projection (UMAP) visualisation showing distinct VSMC subpopulations identified by scRNA-seq of CAD vs control transplant coronary arteries.

To assess whether ox-LDL response genes *in vitro* are modulated in Human coronary artery VSMCs *in vivo*, we meta-analysed VSMCs from coronary artery plaques from people with CAD and from normal coronary arteries from people without CAD (**Fig-S3**). Pseudobulk analysis showed that 2,840 of the 11,225 normalised genes from VSMCs in coronary artery plaques were significantly differentially expressed compared to expression levels in the normal arteries from people without CAD. Of these genes, the expression of 591was altered by ox-LDL in Human coronary artery VSMCs *in vitro*, including 117 that were concordantly upregulated and 119 that were concordantly downregulated in plaques compared to normal arteries and also by ox-LDL *ex vivo*. Examples included, *ANGPTL4* and *NGFR* which were both upregulated in both datasets, and *RASL11A* and *PTGIS* which were both downregulated (**Fig-1F**).

To investigate the differences between VSMCs in CAD and those exposed to ox-LDL *ex vivo*, we performed clustering of the VSMC populations using scRNA-seq data. Previously, Wirka et a^l30^ reported a second ‘fibromyocyte’ cluster with progressive reduction of contractile markers, *ACTA2*, *MYL9*, *CNN1*, *TAGLN*, *TPM2*, but increased expression of modulated VSMC, *IGFBP2*, *BGN*, *FN1*, *GAS6*, *MFGE8*, and *MGP*. Hu et al^31^ reported two additional clusters, synthetic (high in *FN1*, *VCAN*, *S100A10*, *IGFBP6*, and *PRSS23*) and proliferative (high in *MT1A*, *FABP4*, *CCL4*, *MT1M*, and *RGS16*). Our meta-analysis of plaque cells identified six VSMC clusters, including contractile (*CNN1*, *TAGLN*, *TPM2*, *ACTA2*), synthetic with inflammatory signals, pre-foam-like (CD36, FABP4/5) with hypoxia adaptation (*LDHA*, *COX4I2*, *SLC2A3*), fibromyocyte (*COL6A2*, *COL16A1*, *COL4A1*, *THBS2*, *HSPG2*) with lineage genes (*MYOCD*, *PBX1*), fibromyocyte-like with ECM (*DCN*, *FMOD*, *LUM*, *ECM1*, *FBLN1/2/7*), PDGF (*PDGFRB*, *PDGFD*, *NRP1*), complement/immune–matrix (*C3*, *C7*, *FCGRT*), and lipid genes(*ABCA8*), and interferon (IFN)-driven inflammatory VSMCs (*HLA-A/B/C*, *B2M*, *CD74*, *IFNGR2*, *CCL5*, *IL32*, *VCAM1*, *ICAM1*), (**Fig-1G**, **Tables S4**). Interestingly, we found the ox-LDL response genes *ANGPTL4*, *AKR1C1*, *NGFR* to be enriched in cells in pro-inflammatory, foam-like, or synthetic/proliferative-inflammatory states (**Fig-S4**). Comparing plaques to normal artery, we observed significantly higher expression levels of *ANGPTL4* in all clusters and *NGFR* in only synthetic and pre-foam clusters. Conversely, the ox-LDL-downregulated genes *KLF2*, *FOSB*, and *PTGIS* were mainly enriched in cells in contractile, synthetic, or fibromyocyte states.

### Ox-LDL predominantly remodels distal *cis*-regulatory elements

Given the marked ox-LDL-induced changes in gene expression in Human coronary artery VSMCs, we investigated how the underlying enhancer–promoter landscape is remodelled by ox-LDL. We performed ATAC-seq to profile chromatin accessibility and detected ∼169k open chromatin regions (OCRs). Ox-LDL induced significant alterations in the accessibility at 22,081 (13%) of these OCRs (*P*.adj <0.05, **Fig-2A**, **Fig-S5A**), of which 9,659 had increased and 12,422 had decreased accessibilities. Changes in the H3K27ac ChIP-mentation signal triggered by ox-LDL are shown in **Fig-S5B–C**.

**Fig-2.**
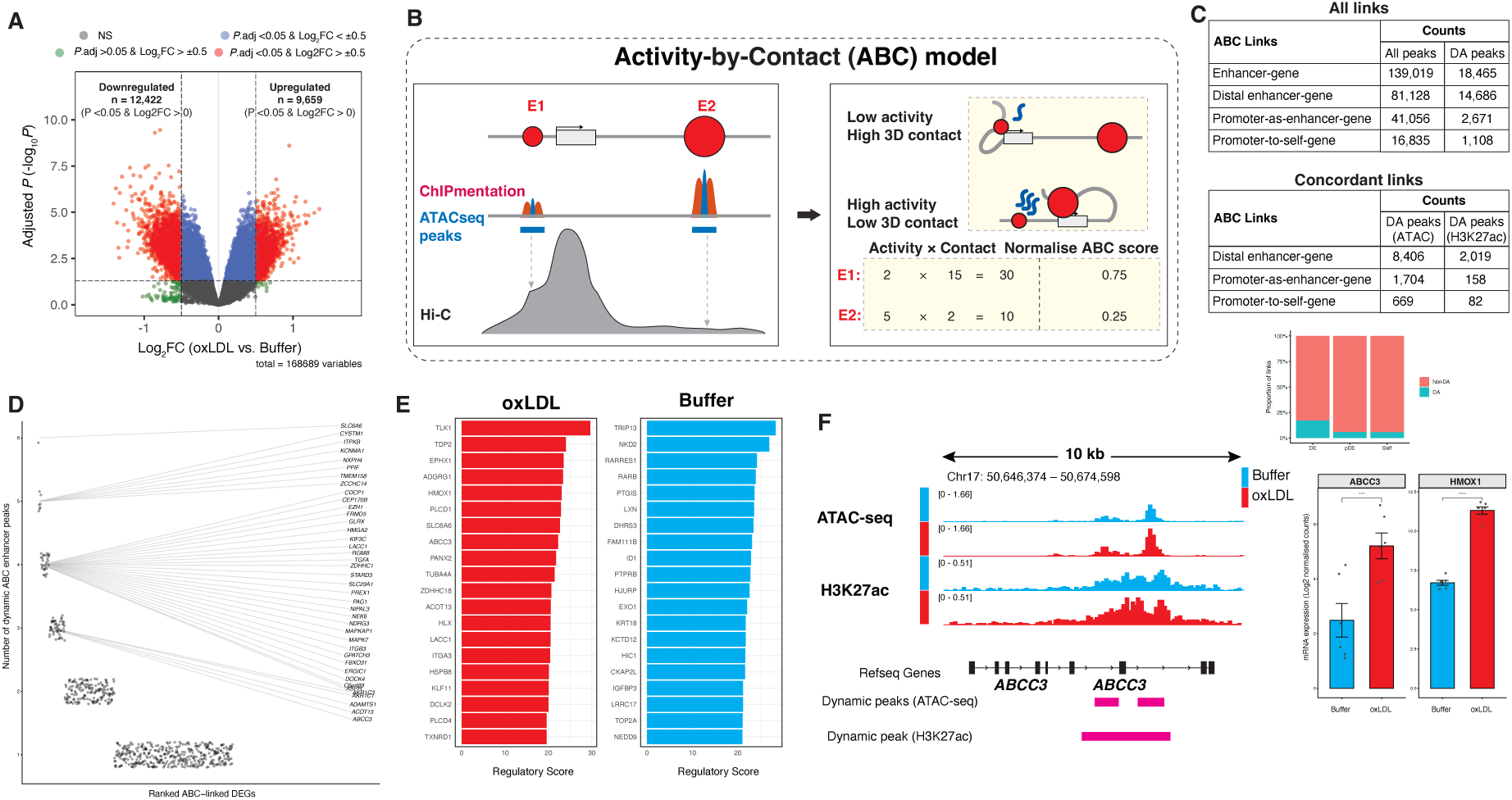
Multi-omics of differentially active enhancers mediating ox-LDL transcriptional regulatory programs in coronary artery VSMCs. **(A)** Volcano plot of the accessible chromatin peaks (ATAC-seq) induced by ox-LDL compared to buffer. The horizontal dashed line represents P.adj <0.05 and vertical dashed lines are log_2_ fold change (log_2_FC) = -0.5 or +0.5. NS, not significant; adjusted P indicates FDR-corrected P-values. **(B)** Illustration of the Activity-by-Contact (ABC) analysis pipeline. **(C)** Summary of the ABC analysis showing predicted enhancer-gene links with ABC score >0.015. DE, distal enhancer; pDE, promoter-as-enhancer; Self, promoter **(D)** Ranking of ox-LDL-upregulated ABC-linked DEGs by number of dynamically active distal enhancers. Each point is an ABC-linked DEG. y-axis: number of dynamic distal enhancers linked to that gene; x-axis: genes ranked by this count (descending). Most genes are linked to a single enhancer, but a subset is predicted to be regulated by multiple enhancers (up to 6 per gene in ox-LDL; Buffer in **Fig-S6B**). **(E)** Top 20 ABC linked DEGs upregulated (Left) and downregulated (Right) by ox-LDL with the highest regulatory scores. **F:** Representative genome browser view illustrating two upregulated intronic enhancers ∼25 kb and ∼27 kb downstream of the TSS of the ATP binding cassette subfamily C member 3 (ABCC3) gene. Expression levels of some of the target DEGs are shown to right.

To map the dynamic OCRs to *cis*-regulatory elements (CREs), we constructed a genome-wide enhancer-gene (E-G) map by integrating data from ATAC-seq (accessibility), H3K27ac ChIPmentation (activity), and Hi-C contact frequencies using the Activity-By-Contact (ABC) model (**Fig-2B**). The ABC map comprised 139,019 enhancer–gene (E–G) links: 81,128 distal enhancer links (median distance 80.7 kb from the target TSS); 41,056 promoter-as-enhancer links, where the promoter of one gene acts as an enhancer for a different gene (median 109.2 kb); and 16,835 promoter-to-self links, in which a promoter contacts its own gene (local promoter–TSS interactions, median 190.5 bp). Thus, promoter-as-enhancer links capture long-range reuse of promoters as distal regulatory elements, whereas promoter-to-self links represent short-range, canonical promoter–gene contacts.

We then overlaid the previously defined differential ATAC-seq peaks onto ABC candidate regions to annotate dynamic versus non-dynamic CREs. Among ATAC-seq peaks overlapping distal ABC enhancers, 17.0% were differentially accessible (∼15k peaks, **Fig-2C**). After restricting E–G links to expressed genes, 24.1% of these differentially accessible distal regions showed concordant H3K27ac changes (edgeR FDR < 0.05; **Fig-2C**). A smaller proportion of promoter-as-enhancer and promoter-to-self classes were differentially accessible (5.3% and 5.1%), with 9.3% and 12.3% of these having concordant H3K27ac signal, respectively. In total, dynamic CREs connect to 2,008 DEGs via 4,243 peak–gene links, indicating that ox-LDL-driven chromatin remodelling is concentrated at distal regulatory elements and mechanistically linked to differential gene expression.

We then focused on concordant ABC enhancer-gene links (i.e., links where changes in accessibility are correlated with concordant changes in gene expression). Most dynamic distal enhancers had one-to-one links with target DEGs, but some were pleiotropic; for example, 11 ox-LDL-responsive distal enhancers were each predicted to regulate three DEGs (*P*.adj <0.05, **Fig-S6A**). In turn, several ABC-linked DEGs were predicted to be regulated by multiple dynamic distal enhancers, up to six per DEG in ox-LDL-enriched (**Fig-2D**) and up to nine per DEG in repressed enhancers (**Fig-S6B**). These patterns are consistent with coordinated enhancer–gene regulation, and this is supported by a significant positive correlation between enhancer accessibility and H3K27ac signal across concordant enhancer-gene links (N = 886, R = 0.43, *P* <0.001, **Fig-S6C**). Distal enhancers linked to ox-LDL-upregulated ABC-linked DEGs showed higher accessibility than enhancers linked to non-DEGs or to ox-LDL-downregulated DEGs (*P* <0.001, **Fig-S6D**). The top 20 highly concordant ABC-linked DEGs are shown in **Fig-2D** and **2E**, including *ABCC3*, *HMOX1*, *TLK1*, *KLF11*. Finally, pathway enrichment of all genes ABC-linked to dynamically upregulated distal enhancers highlighted oxidative stress, autophagy, apoptosis, muscle differentiation, smooth muscle migration, regulation of hypoxia, mirroring pathway enrichment of transcriptomics signatures (*P*.adj <0.05, **Table S5**). In summary, these data show that ox-LDL reprograms HCASMC enhancer–gene networks towards oxidative stress, autophagy, apoptotic, and migratory transcriptional programmes.

### Genome-wide footprinting reveals ox-LDL-induced transcription factor rewiring

The multiomics analysis demonstrates that ox-LDL induces changes in chromatin accessibility and H3K27ac signal that are mechanistically associated with changes in gene expression through chromatin contacts. We hypothesised that the chromatin changes alter transcription factor (TF) binding. To test this, we first performed genome-wide TF footprinting on ATAC-seq profiles using TOBIAS to infer changes in TF motif occupancy resulting from exposure of Human coronary artery VSMCs to ox-LDL. Ox-LDL led to significant changes in TOBIAS motif occupancy scores at 330 motifs (*P*.adj <0.05 & logFC ≥0.01) corresponding to 305 unique TFs that were expressed in Human coronary artery VSMCs (by RNA-seq), of which 96 were differentially expressed in response to ox-LDL (RNA-seq *P*.adj <0.05; **Fig-3A**, **Fig-S7**). The number of significant motifs exceeds the number of TFs because several TFs share closely related motif variants that are represented as distinct motif models. We then prioritised TFs whose motifs showed both significant changes in inferred occupancy and significant differential expression (RNA-seq <0.05). This identified 24 TFs (27 motifs) with increased motif occupancy in ox-LDL-treated cells (ox-LDL footprints) and 40 TFs (48 motifs) with reduced occupancy in ox-LDL-treated cells. Among the motifs with the largest increases in inferred occupancy were a MEF2D cluster (MEF2A & MEF2D), ESRRA, and MSC with the largest reductions in inferred occupancy being for GATA6, a FOXO2 cluster (FOXO1, FOXO2), and a KLF5 cluster (KLF2, KLF5) (**Fig-3B–3C**).

**Fig-3.**
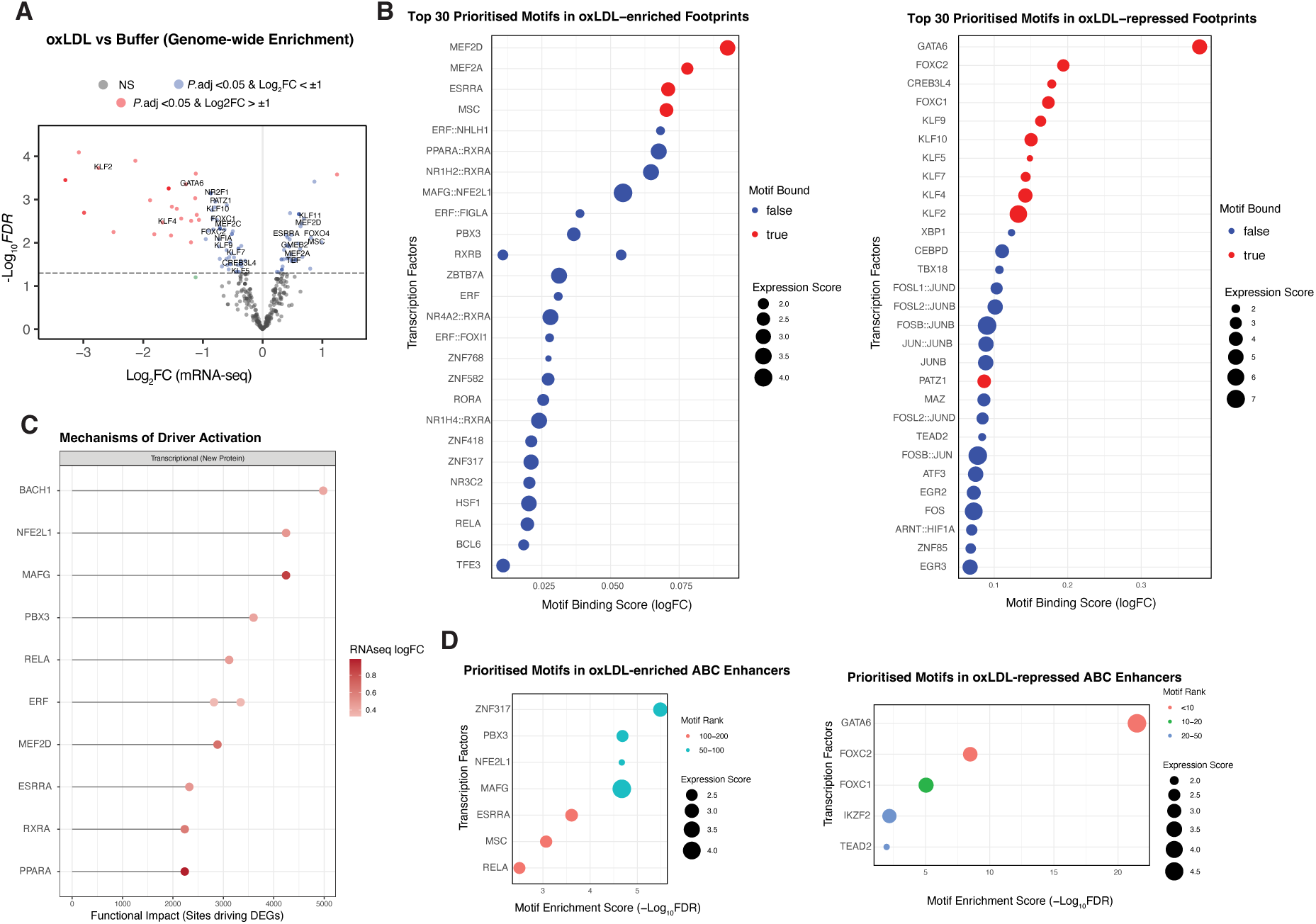
Genome-wide footprinting and motif analysis identify transcriptional regulators of the ox-LDL response. **(A)** Volcano plot of differential TF motif occupancy inferred from ATAC-seq footprints (TOBIAS) comparing ox-LDL vs Buffer. x-axis: log_2_ fold-change in TF expression in RNA-seq (ox-LDL–Buffer); y-axis: −log_10_(FDR). Significant motifs (FDR < 0.05; èect threshold as in Methods) are annotated to expressed TFs; labels highlight TFs that are both differentially bound and RNA-seq DEGs (TMM-normalised; see Methods). **(B)** Representative concordant TFs: examples with increased (LEFT) or reduced (RIGHT) binding/expression following ox-LDL exposure. Concordant = same directional change in footprinting and transcript abundance. **(C)** Prioritised TFs at ABC enhancers linked to DEGs that show coordinated activation with ox-LDL, defined by increased TF expression (RNA-seq) and increased motif accessibility (positive chromVAR differential deviation). TFs are ranked by the number of ABC-linked enhancer sites assigned to DEG targets and coloured by RNA-seq logFC. Motif accessibility was quantified by chromVAR, TF–target relationships were inferred by integrating TF motif sites with ABC enhancer–gene links, and TFs were further restricted to those supported by TOBIAS footprinting. **(D)** Motif enrichment changes among ABC-linked distal enhancers overlapping dynamic CREs using AME (MEME Suite) with length/GC-matched background; FDR < 0.05. Displayed TFs pass the concordance filter (DEG direction matches CRE direction). ABC parameters and promoter analyses are detailed in Methods and **Fig-S8**; expanded TF lists in **Fig-S7**.

Next, we analysed TF motif enrichment on dynamic CRE peaks linked to ABC enhancers using Analysis of Motif Enrichment of MEME-SUITEs. We performed enrichment analysis of ox-LDL-upregulated and downregulated peaks in enhancer CREs. This identified 83 motifs bound by 74 unique TFs in distal enhancers and 19 motifs bound by 14 unique TFs in promoters (**Fig-S8**). Applying a concordance filter (ox-LDL-upregulated DEGs ABC-linked to ox-LDL-upregulated CREs; downregulated DEGs ABC-linked to ox-LDL-downregulated CREs) yielded 7 TFs in ox-LDL-enriched enhancers, 3 TFs in ox-LDL-enriched promoters, and 5 TFs in ox-LDL-repressed enhancers (**Fig-3D, Fig-S8C**). Transcription factors that are enriched in both footprinting and enhancer analyses included C2H2 ZNFs (>3 zinc fingers, e.g., ZNF317), bZIPs (BACH1, MAFG), MSC, PBX3, and RELA. Conversely, ox-LDL-repressed regulatory programmes were associated with motifs for GATA6, FOXC1, FOXC2, and TEAD2.

Overall, ox-LDL shifts VSMC gene regulatory programme from a FOXO/KLF/GATA-centric homeostatic signatures toward MEF2, ESRRA, MSC, and bZIP-family signatures together with NF-κB pathway-associated transcriptional activity, with concordant signals across genome-wide footprints and ABC-linked enhancer motif enrichment.

### CAD risk variants are enriched in ox-LDL-responsive enhancers and converge on target genes

Several studies have shown that genetic variants within GWAS loci often function in a tissue- or cell type-specific manner.^11^ We hypothesised that disease-associated CAD variants would modulate ox-LDL response in coronary VSMCs, and that these effects would be context-dependent rather than uniformly observed across cell types. To test this, we defined a comprehensive set of CAD risk variants by pruning the 393 genome-wide significant CAD loci in the NHGRI-EBI GWAS Catalog (*P* < 5×10^-8^) based on LD to obtain approximately independent index SNPs. Each index SNP was then expanded to all common dbSNP v155 variants (minor allele frequency > 1%) in strong LD (r^2^ ≥ 0.8, EUR), yielding a final panel of 19,335 CAD-associated variants for downstream enrichment analyses (**Fig-4A**).

**Fig-4.**
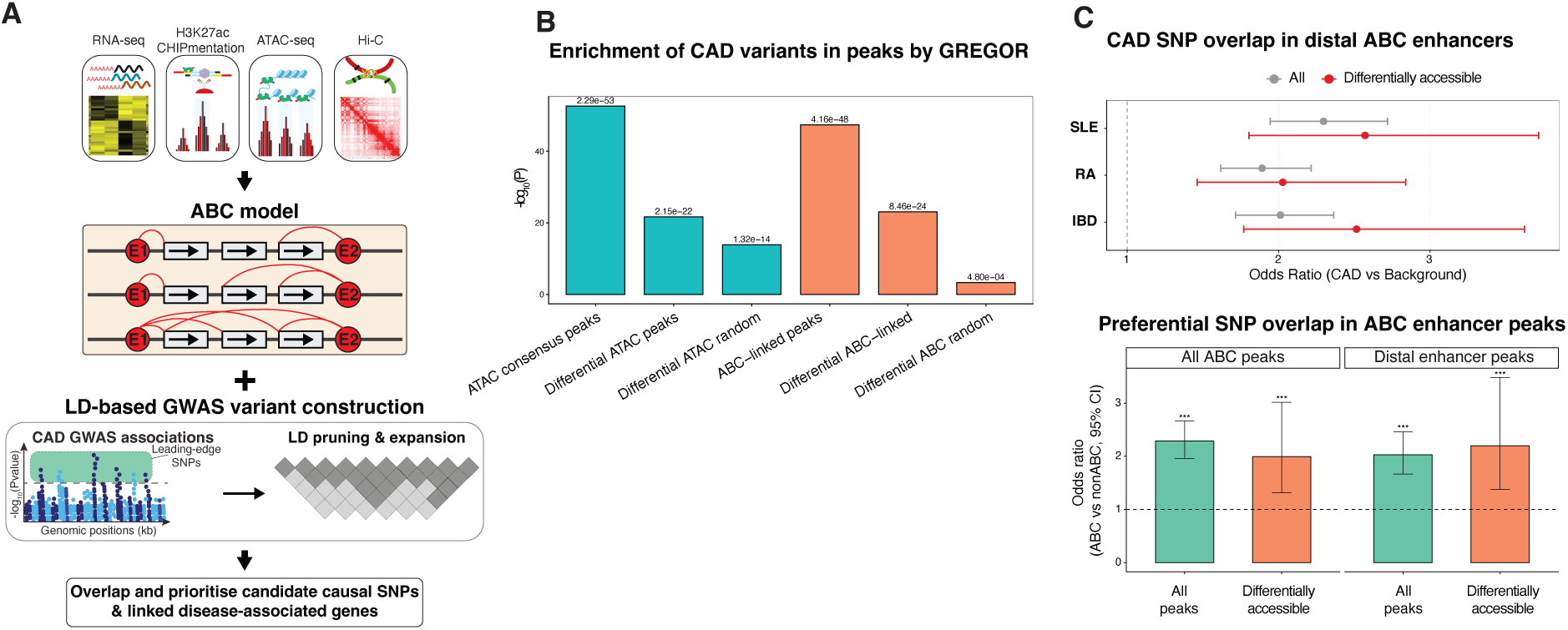
Integration of multiomics data with GWAS prioritises CAD candidate causal genes mediating the ox-LDL response in coronary artery VSMCs. **(A)** Genetic analysis framework. CAD GWAS single-nucleotide polymorphisms (SNPs) were downloaded from the EMBL–EBI GWAS Catalog and filtered to retain variants reaching genome-wide significance (P < 5.0 × 10^-8^). These SNPs were pruned based on high linkage disequilibrium (LD) to obtain approximately independent index SNPs. Each index SNP was expanded to all common dbSNP v155 variants (MAF > 1%) in strong LD (r^2^ ≥ 0.8, EUR), yielding a final panel of 19,335 CAD-associated variants. Resulting SNPs were intersected with dynamic ABC candidate regulatory element (CRE) peaks, and candidates were prioritised if they were linked to differentially expressed genes (DEGs). **(B)** GREGOR analysis of CAD GWAS SNP enrichment in regulatory annotations. Bars show significance as −log_10_(P) for overlap of CAD index SNPs and their LD proxies within all ATAC-seq peaks, differential ATAC-seq peaks, all ABC-linked peaks, and differential ABC-linked peaks. Expected overlap was estimated using matched control SNP sets selected by GREGOR to account for LD structure, minor allele frequency, and distance to the nearest gene. P-values were calculated by GREGOR from the observed versus expected overlap. Random size-matched genomic regions were included as negative controls. **(C) Top:** Overlap analysis results for ABC distal enhancers comparing CAD variants relative to control variants associated with specific autoimmune diseases. Odds ratios are shown for both the full set (All) and the subset of differentially accessible distal enhancers to demonstrate CAD-specific signalling. IBD, inflammatory bowel disease; SLE, systemic lupus erythematosus; and RA, rheumatoid arthritis. **Bottom:** Overlap of CAD variants in ATAC-seq peaks linked to target genes (ABC peaks) compared to unlinked peaks (non-ABC). This comparison is presented for all ABC peaks and for the subset of differentially accessible ABC peaks.

We first used GREGOR to assess whether CAD GWAS index SNPs were enriched within accessible chromatin and ABC-linked regulatory regions. All ATAC-seq consensus peaks were significantly enriched for CAD SNPs (603 index SNPs overlapping vs 388 expected; *P* = 2.29 × 10^-53^), and this enrichment was maintained in differentially accessible peaks (193 vs 100 expected; *P* = 2.15 × 10^-22^). Similarly, all ABC-linked peaks showed significant CAD SNP enrichment (372 vs 194 expected; *P* = 4.16 × 10^-48^), with differential ABC-linked peaks retaining strong enrichment (118 vs 46 expected; P = 8.46 × 10^-24^). Size-matched random genomic regions showed markedly attenuated or no enrichment (e.g., differential ABC random sets: P = 4.8 × 10^-4^ and P = 0.35), confirming that the observed enrichment reflects concentration of CAD risk variants within biologically active, ox-LDL-responsive regulatory elements.

We next compared the overlap of CAD variants in CREs relative to variants associated with immune disease controls (IBD, SLE, RA). Within distal enhancers, CAD variants showed significant overlap compared to control variants (odds ratio [OR] were 2.01, 2.30 and 1.89 for CAD compared to IBD, SLE and RA respectively **Fig-4B**), with an even stronger overlap for the subset of variants in dynamically accessible chromatin (IBD: OR 2.51, SLE: OR 2.57, RA: OR 2.03). In contrast, promoter-distal elements and self-loops did not show significant overlap in the differentially accessible subset (**Fig-S9**), consistent with the distinct regulatory roles of dynamic enhancers compared to constitutive promoters. Consistently, at the peak level, ABC-linked peaks were more likely than non-ABC peaks to harbor CAD SNPs (All peaks: OR 1.99 [1.31–3.02], *P* <0.01; differentially accessible peaks: OR 2.20 [1.38–3.48], *P* <0.01), indicating that ABC-mapped enhancers capture a CAD-enriched subset of the ox-LDL-responsive regulatory landscape.

Peaks with ox-LDL-induced differential accessibility overlapped with 75 SNPs in 51 CAD loci (**Table S6**), including 66 SNPs (47 loci) in ABC distal enhancers. To understand the potential impact of these CAD SNPs in ox-LDL-modulated enhancers, we studied their ABC target genes. In total, CAD-associated SNPs in dynamic enhancers were predicted to regulate 83 target genes, with an average of 1.86 SNPs-to-gene links. Of these, 32 SNPs at 22 CAD loci were associated with differential expression of 28 genes, including 10 that were upregulated and 18 that were downregulated in response to ox-LDL. Enhancers that had increased chromatin accessibility with ox-LDL harboured 23 CAD SNPs and were ABC-linked to 20 differentially expressed genes, with *MAP1S*, *WBP2*, *SLC44A2*, *IL6R*, *DCLK2*, and *UBAP2L* having the highest concordance between ox-LDL-mediated enhancer (ABC score) and target gene expression (**Fig-5A&B**). Conversely, those that had reduced chromatin accessibility with ox-LDL harbored nine CAD SNPs and were ABC-linked to 9 differentially expressed genes, of which eight were ox-LDL-downregulated genes, (**Fig-6A**).

**Fig-5.**
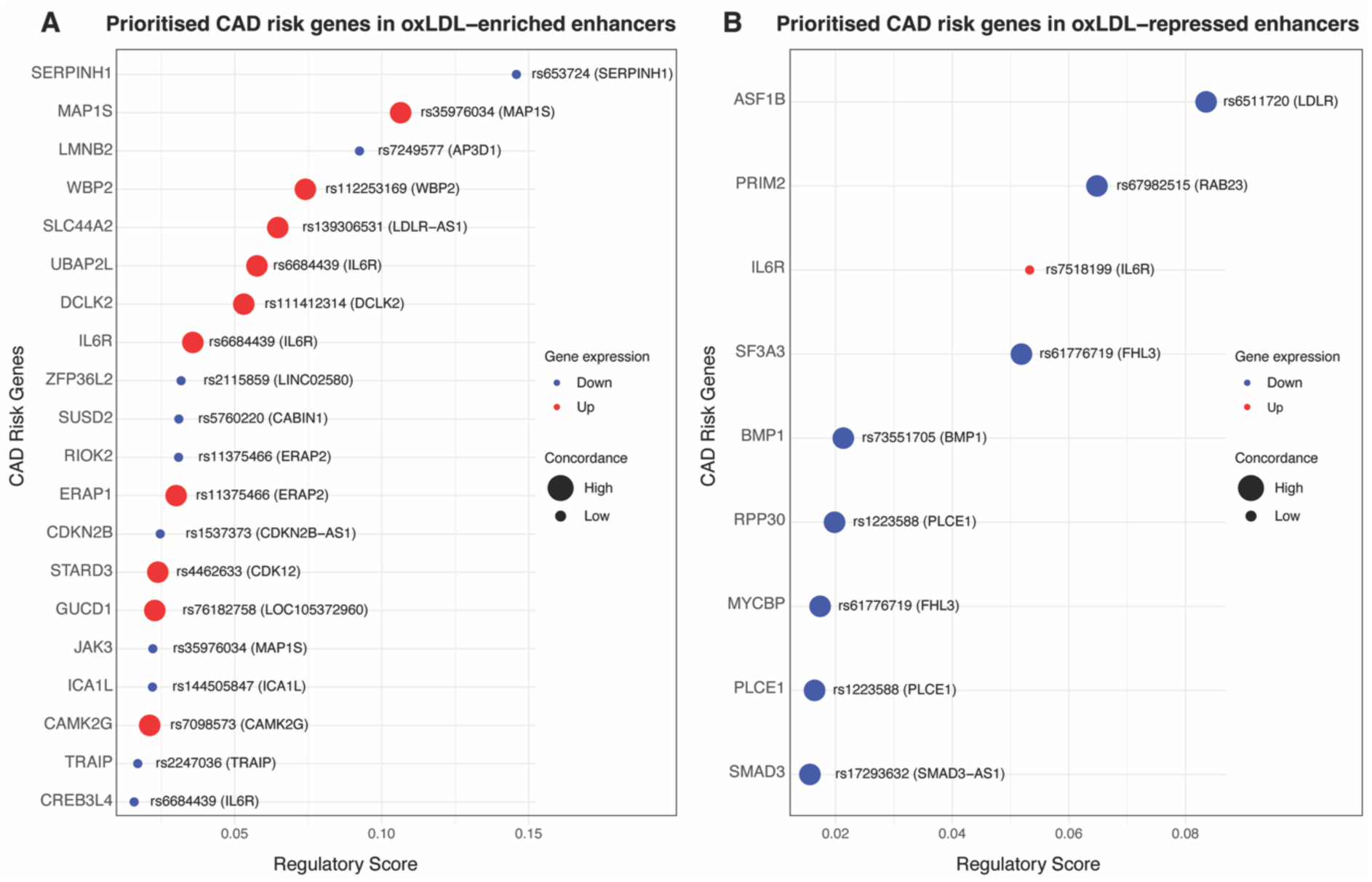
Prioritised CAD SNPs in ox-LDL modulated enhancers in Human coronary artery VSMCs. **(A)** Top ox-LDL-upregulated genes linked by ABC to distal enhancers that have increased chromatin accessibility with ox-LDL and overlap CAD-associated SNPs. **(B)** Top ox-LDL-downregulated genes linked by ABC to distal enhancers that have reduced chromatin accessibility with ox-LDL and overlap CAD-associated SNPs. SNP IDs and their loci (in parentheses) are shown. The regulatory score denotes the ABC score for each enhancer–gene pair. Concordance reflects agreement between enhancer and gene responses: high concordance indicates that ox-LDL increases enhancer accessibility and upregulates the linked gene, or decreases enhancer accessibility and downregulates the linked gene; low concordance indicates discordant responses (e.g., increased accessibility with gene downregulation, or decreased accessibility with gene upregulation).

Together, these data show that CAD risk loci appear to influence disease by perturbing ox-LDL-responsive distal enhancer–gene interactions in coronary artery VSMCs.

We next generated CAD credible sets to prioritise variants with high posterior probability of being causal at each GWAS locus. Credible sets were derived by statistical fine-mapping of CAD summary statistics (SuSiE-rss) using an external UKBB EUR LD reference to define 95% credible sets. Overlaying these credible sets onto the ABC enhancer map showed that peaks with ox-LDL-induced differential accessibility overlapped 20 credible-set SNPs across 16 CAD loci (**Table S7**), including 19 SNPs (15 loci) within ABC-linked distal enhancers. These fine-mapped CAD variants in dynamic enhancers were linked to 35 target genes, with an average of 2.74 SNP–enhancer–gene links per SNP. The credible-set analysis further prioritised two additional DEGs, *ST3GAL4* and *HIC1* (**Fig-S9**). This shows that intersecting fine-mapped CAD credible sets with ox-LDL-responsive ABC enhancers refines the genetic signal to a small number of high-confidence SNP–enhancer–gene pairs likely to mediate CAD risk.

Taken together, these analyses show that CAD risk variants, including fine-mapped credible-set SNPs, are concentrated in ox-LDL-responsive distal ABC enhancers that converge on a focused set of target genes, providing a mechanistic link between CAD loci and disease-relevant transcriptional programs in Human coronary artery VSMCs

### Allele-specific modelling nominates TF-mediated enhancer mechanisms at CAD variants

Given the enrichment of CAD risk variants within ox-LDL-responsive regulatory programmes, we next interrogated putative mechanisms at the prioritised loci using orthogonal *in-silico* approaches. First, we applied AlphaGenome to predict allele-specific regulatory effects on chromatin activity (accessibility and histone modifications) and transcriptional output (RNA-seq and CAGE-seq) across a 1 Mb sequence context in vascular smooth muscle cells (**Fig-6A**). Relative to a non-vascular comparator (B cells), AlphaGenome predicted stronger effects of prioritised CAD variants on active chromatin features in vascular smooth muscle cells, including H3K27ac, H3K4me1/2/3 and H3K9ac (**Fig-6B**). These predicted shifts were accompanied by larger inferred impacts of disease-associated variants on transcriptional readouts, including RNA-seq signal and the active gene-body mark H3K79me2 in VSMC compared to B cells; in contrast, predicted effects on the repressive mark H3K27me3 were reduced in VSMC compared with B cells.

**Fig-6.**
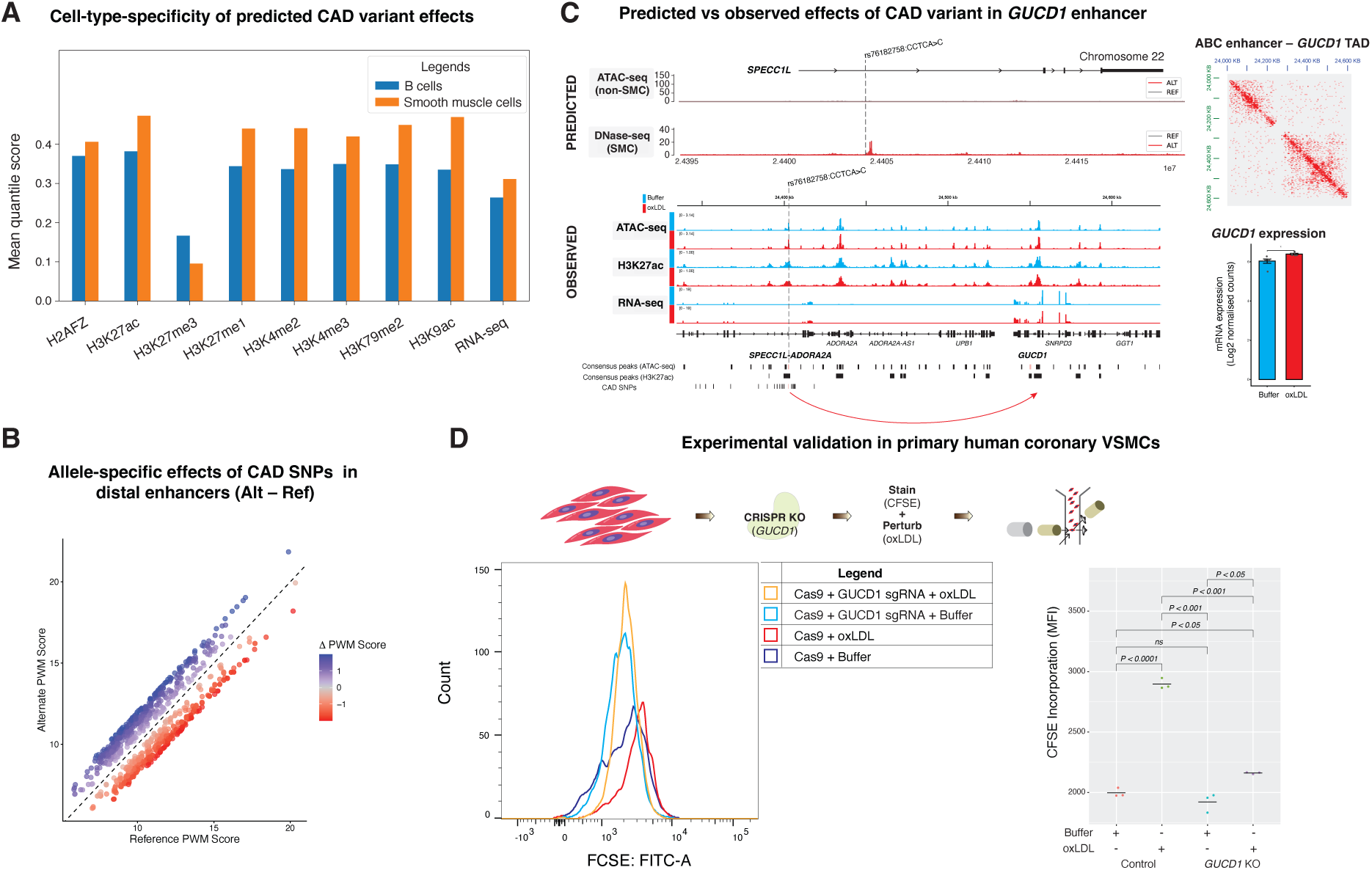
Mechanisms of CAD risk variants in ox-LDL-responsive enhancers. **(A)** AlphaGenome predicted quantile scores for prioritised CAD variants in vascular smooth muscle cells versus a non-vascular comparator (B cells) across chromatin and transcriptional output tracks. The quantile score represents a standardised rank of the impact of a variant, derived from quantile normalisation of the raw predicted score for each variant. **(B)** In-silico motif disruption analysis (motifbreakR) for prioritised variants within ABC enhancers, showing reference versus alternate PWM scores, coloured by ΔPWM (alternate − reference). TFs were restricted to those expressed in Human coronary artery VSMCs. **(C)** Locus-level example at rs9624456:CCTCA>T. Top left: AlphaGenome-predicted genome tracks comparing alternate versus reference alleles at rs9624456. Bottom left: observed genome tracks in Human coronary artery VSMCs following ox-LDL exposure. Top right: Hi-C contact map showing chromatin interactions within the SPECC1L-GUCD1 topologically associating domain (TAD). Bottom right: RNA-seq shows ox-LDL-associated induction of GUCD1 in Human coronary artery VSMCs. **(D)** CRISPR knockout (KO) experimental design and proliferation readout. Human coronary artery VSMCs were transfected with sgRNA, labelled with CFSE, and exposed to ox-LDL. CFSE histograms and quantification are shown; higher CFSE signal indicates reduced proliferation.

Second, we assessed allele-dependent TF motif perturbation using motifbreakR, restricting analyses to TF expressed in Human coronary artery VSMCs. Among prioritised variants residing in ABC enhancers, alternate alleles showed systematically higher motif scores than reference alleles, yielding positive ΔPWM values across sites (**Fig-6C, Fig-S10A**). Together, these analyses support a model in which CAD variants in ox-LDL-responsive ABC enhancers act in a cell-type-specific manner, preferentially modulating active regulatory chromatin and TF binding potential in Human coronary artery VSMCs.

### *SPECC1L* locus maps to *GUCD1* and modulates ox-LDL-linked coronary VSMC proliferation and senescence

At the *SPECC1L-ADORA2A* locus, we prioritised rs76182758 because it lies within an active ox-LDL-responsive enhancer that became more accessible following ox-LDL exposure (ATAC FDR = 0.006). This variant is in high LD with rs9624456 (r^2^ > 0.8), which has been associated with CAD in a trans-ancestry GWAS meta-analysis of Japanese and European populations. To interrogate allele-specific mechanism, we used AlphaGenome to predict regulatory effects of the alternate ‘C’ versus reference ‘CCTCA’ allele. AlphaGenome predicted a vascular smooth muscle cell–restricted increase in DNase signal for the ‘C’ allele, with no corresponding effect in non-vascular cells (**Fig-6D**). Consistent with these predictions, ox-LDL exposure increased accessibility at this enhancer in human coronary artery VSMCs.

To interrogate the allele-specific mechanism, we used AlphaGenome to predict regulatory effects of the alternate ‘C’ versus reference ‘CCTCA’ allele. In both AlphaGenome predictions and experimental data on the response to ox-LDL, we found no evidence that rs76182758 influences local *SPECC1L* expression, suggesting a distal regulatory target. Using the ABC model, we linked the enhancer harbouring rs76182758 to *GUCD1* (∼120 kb downstream), supported by a high contact frequency between the enhancer and *GUCD1* in Hi-C maps (**Fig-6D**, top right). In keeping with enhancer activation, ox-LDL exposure increased *GUCD1* expression (**Fig-6D**, bottom right), providing convergent evidence that this CAD locus has an ox-LDL-induced regulatory effect on *GUCD1* rather than *SPECC1L*.

To test the potential functional relevance of *GUCD1* in the response to ox-LDL, we performed CRISPR-based knockout in human coronary artery VSMCs using an sgRNA targeted to exon 3 of GUCD1. When a control sgRNA was used, ox-LDL significantly inhibited proliferation (**Fig-6E**; P < 0.001). In contrast, with the *GUCD1-*targetting sgRNA knockout the inhibitory effect of ox-LDL on proliferation was abolished and proliferation was restored towards that of relative to ox-LDL-treated controls (**Fig-6D**; *GUCD1* sgRNA/ox-LDL vs control /ox-LDL, P < 0.001). *GUCD1* knockout also reduced expression of the senescent marker p21 (**Fig-S11**, *P* < 0.05)

Together, these data nominate *GUCD1* as the effector gene of the enhancer variant rs76182758 at the *SPECC1L* locus and place this axis in a pathway linking CAD genetic risk to HCASMC proliferative/senescent responses following ox-LDL exposure.

### *MAP1S* locus modulates proliferation through a *BACH1*–*MAP1S* axis

At the *MAP1S* locus, we prioritised the variant rs35976034 due to its location within an intronic enhancer that had increased chromatin accessibility following ox-LDL exposure (ATAC-seq FDR = 0.04). This SNP is in high LD (r^2^ > 0.8) with the lead CAD signal rs10410487. AlphaGenome predicted allele-specific regulatory activity at this locus, with the alternate ‘T’ allele associated with increased *MAP1S* activation relative to the reference ‘G’ allele, supported by higher predicted H3K4me3 and CAGE-seq signal (**Fig-7A**). Experimental data is consistent with this as ox-LDL exposure led to chromatin hyper-accessibility at the enhancer in human coronary artery VSMCs (**Fig-7A**).

**Fig-7.**
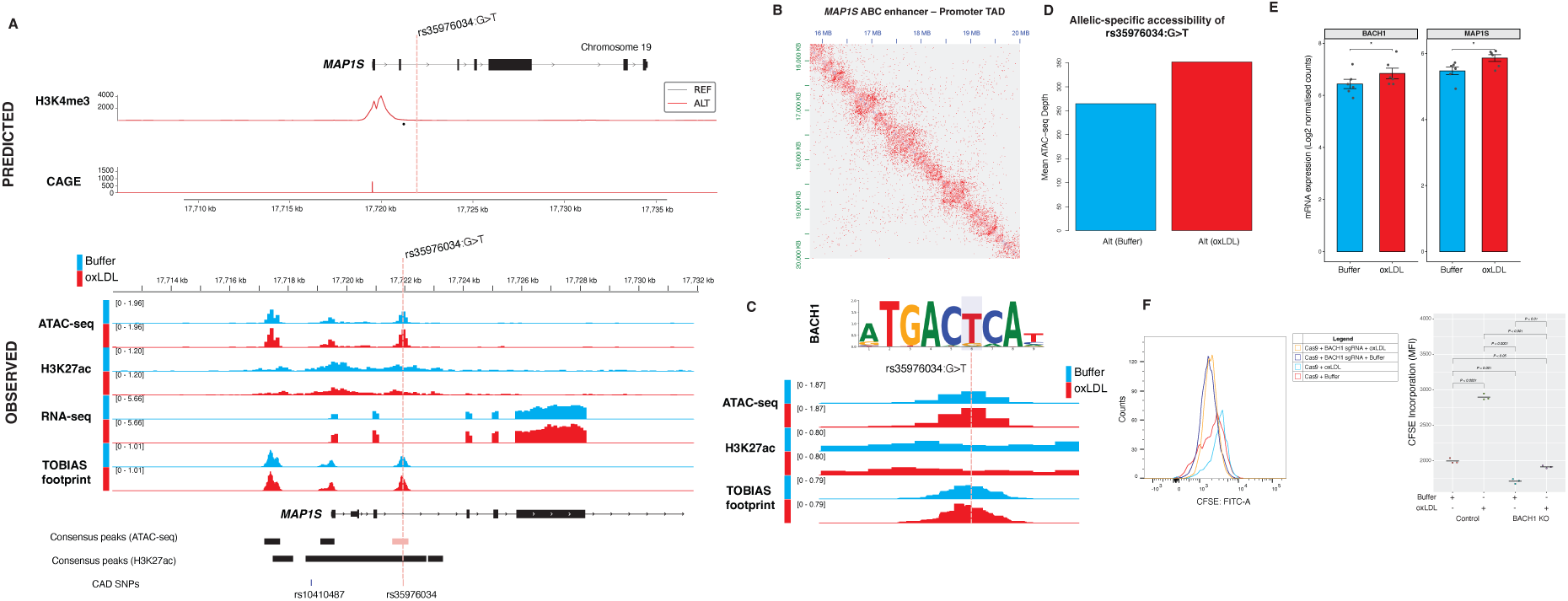
Mechanism of variant in an ox-LDL-responsive enhancer at the MAP1S locus. **(A)** Locus-level example at rs35976034 (G>T). **Top:** AlphaGenome-predicted genome tracks comparing alternate versus reference alleles. **Bottom:** observed genome tracks in Human coronary artery VSMCs following ox-LDL exposure (b. **(B)** Hi-C contact map showing chromatin interactions within the locus TAD encompassing the enhancer and the MAP1S promoter. **(C)** Higher resolution coverage plots centred on rs35976034, with prioritised BACH1 motif shown at the top. **(D)** Allele-specific ATAC-seq signal at rs35976034, showing mean read depth for the alternate ‘T’ allele under buffer and ox-LDL conditions. **(E)** RNA-seq shows ox-LDL-associated inductions of MAP1S and BACH1 in Human coronary artery VSMCs. **(F)** CRISPR KO and CFSE-based proliferation readout under buffer and ox-LDL conditions.

ABC modelling linked the rs35976034-containing enhancer to the *MAP1S* promoter, supporting direct enhancer–promoter coupling within the locus (**Fig-7A, B**). To nominate trans-acting regulators, TOBIAS footprinting demonstrated that exposure to ox-LDL increased TF occupancy at the enhancer, and variant-to-motif mapping implicated *BACH1* as a candidate regulator at the rs35976034 site (**Fig-7C**). This was supported by a global ox-LDL-induced increase in *BACH1* motif accessibility (chromVAR deviation = 4.08). Descriptive allele-specific read-depth analysis at rs35976034 in a heterozygous donor is shown in **Fig-S10B**.

At the transcript level, ox-LDL significantly increased both *MAP1S* and *BACH1* expression (**Fig-7E**), supporting a coherent regulatory model in which ox-LDL induces *BACH1* activity at the rs35976034 enhancer to promote *MAP1S* expression. Functionally, CRISPR knockout of *BACH1* in Human coronary artery VSMCs reversed the ox-LDL-induced inhibition of proliferation seen with control sgRNA (**Fig-7F**; P < 0.001) and reduces cellular senescence markers (**Fig-S12**). Collectively, these data prioritise a *BACH1*–*MAP1S* effector axis at this locus and support rs35976034 as a functional variant within an ox-LDL-responsive enhancer that shapes HCASMC proliferation/senescence responses following ox-LDL exposure.

## DISCUSSION

### Major Findings

Vascular SMCs are central to atherosclerosis, yet how ox-LDL influences coronary VSMC gene regulation has remained unclear. Using multiomic profiling in Human coronary artery VSMCs and integrating ABC enhancer–gene maps with CAD genetics, we show that ox-LDL activates CAD-relevant, foam cell-like transcriptional programmes and drives predominantly distal enhancer remodelling linked to differential gene expression. CAD risk variants are enriched in these ox-LDL-responsive distal enhancers. ABC linking showed that these variant-containing enhancers converge on a focused set of target genes. Allele-specific predictions and locus-level validation indicates that there is CAD risk in coronary VSMCs that is mediated by changes in enhancer accessibility/activity and TF-binding that modulate effector gene expression under lipid stress.

### Ox-LDL exposure recapitulates coronary VSMC transcriptional modulation in CAD

Ox-LDL has been implicated in VSMC dysfunction in atherosclerosis, including oxidative stress, the development of foam cell–like features, apoptosis, and cellular senescence.^53-56^ However, prior transcriptional studies of ox-LDL-treated VSMCs have typically focused on selected pathways and/or limited gene sets. For example, microarray profiling reported modulation of 218 and 833 genes in human coronary VSMCs after 3 h and 21 h of ox-LDL exposure, respectively.^57^ Using unbiased bulk RNA-seq, we observed a broader transcriptional response, with ox-LDL exposure altering expression of >3,000 genes. Ox-LDL upregulated genes in lipid-handling and stress-adaptation programmes (oxidative stress, autophagy, hypoxia, and ferroptosis) alongside repression of genes in extracellular matrix (ECM) organisation and cell-cycle/proliferative programmes. To place these *ex-vivo* responses in disease context, we performed a meta-analysis of transcriptomic profiles from coronary artery plaques (CAD) versus non-CAD coronary tissues. We found that ox-LDL-induced genes were preferentially enriched in plaque VSMC clusters exhibiting pro-inflammatory and foam cell-like programmes, whereas ox-LDL-repressed genes mapped to clusters consistent with contractile and fibromyocyte-like states. Given that loss of ECM programmes and acquisition of inflammatory secretory features in lesion cores has been linked to features of plaque vulnerability,^58^ our data are consistent with an ox-LDL-driven shift towards plaque-destabilising VSMC states. Together, these data suggest that ox-LDL shifts coronary VSMCs away from structural/homeostatic programmes and towards stress-response states that may contribute to maladaptive remodelling in plaques. We next asked how this transcriptional switch is encoded in *cis*-regulatory elements (CREs).

### Ox-LDL drives coronary VSMCs towards stress-adaptation programmes

To address this, we constructed a genome-wide enhancer–gene map in Human coronary artery VSMCs using the ABC framework. The ABC model integrates enhancer activity (ATAC-seq and H3K27ac) with physical contact (Hi-C). This reduces reliance on nearest-gene assignment and allows us to interpret ox-LDL responses in terms of specific enhancer–gene units. We found that ox-LDL-driven chromatin remodelling was dominated by distal enhancers, which had concordant changes in accessibility and H3K27ac signals. 4,243 peak–gene links connected dynamic CREs to 2,008 genes which were differentially expressed following ox-LDL exposure. Enhancers that gained accessibility and H3K27ac signal with ox-LDL exposure were preferentially linked to ox-LDL-upregulated genes. Conversely, those that lost accessibility and H3K27ac signal were preferentially linked to ox-LDL-downregulated genes.

Our analysis clarifies which programmes are regulated by ox-LDL-responsive enhancers. Genes linked to ox-LDL-activated distal enhancers were enriched for oxidative stress, autophagy, apoptosis, muscle differentiation, smooth muscle migration and regulation of hypoxia. This indicates that ox-LDL preferentially remodels enhancers controlling stress-adaptation and phenotypic switching in VSMCs. It provides a *cis*-regulatory mechanism for the ox-LDL-induced gene expression changes. It also enables variant-to-enhancer-to-gene assignment for non-coding CAD GWAS signals within oxLDL-responsive regulatory elements.

### Enhancer-gene maps links non-coding CAD variant to target genes in coronary VSMCs

Most CAD risk variants are non-coding and typically fall in intronic or distal intergenic regions. This makes effector-gene assignment using proximity alone unreliable. Large-scale GWAS have reported hundreds of CAD index signals with thousands of linked variants.^11^ We therefore integrated ATAC-seq, H3K27ac ChIPmentation, and Hi-C with the ABC model to map ox-LDL-responsive enhancers to their candidate target genes in Human coronary artery VSMCs. ABC-linked enhancers were significantly enriched for CAD variants relative to non-linked enhancers. The enrichment was strongest in enhancers whose accessibility was altered following ox-LDL exposure. Moreover, this enrichment was selective for CAD variants compared with variants associated with non-vascular inflammatory traits. These results support disease- and cell-context specificity. Consistent with this, sequence-based modelling predicted larger regulatory effects of prioritised CAD variants in smooth muscle cells than in a non-relevant comparator (B cells). These results demonstrate the value of cell-type-resolved maps for variant-to-gene assignment at CAD loci.

### Locus-level analyses nominated *GUCD1* and *MAP1S–BACH1* circuits as plausible mediators of lipid-stress responses

Using this framework, we focused on two loci where multiple data layers converged on specific enhancer–gene mechanisms. At the *SPECC1L-ADORA2A*-tagged CAD locus, we prioritised rs76182758 and implicated *GUCD1* (guanylyl cyclase–containing domain 1) as an ox-LDL-responsive downstream effector gene in Human coronary artery VSMCs. Prior annotation at this locus has often emphasised *SPECC1L* as the gene of interest based on proximity.^10^ Our data instead support distal enhancer-mediated regulation of *GUCD1*. Ox-LDL increased chromatin accessibility at the rs76182758 CRE which was linked by ABC to *GUCD1. GUCD1* expression increased concordantly with ox-LDL exposure. *GUCD1* knockout attenuated ox-LDL-induced growth arrest and senescence phenotypes, supporting functional involvement in the ox-LDL response. Indeed, previous co-expression analyses also place *GUCD1* in inflammation, cell death, and immune signalling programmes, suggesting broader involvement in stress–inflammatory circuitry.^59^ AlphaGenome predicted allele-specific increases in regulatory activity at this locus; this is consistent with the experimentally observed enhancer activation and *GUCD1* upregulation and provides orthogonal support for an allele-dependent mechanism. Together, these findings shift this CAD signal from proximity-based annotation to an oxLDL-responsive enhancer–GUCD1 mechanism and support *GUCD1* as the most plausible effector under lipid-stress in coronary artery VSMCs.

We also identify a MAP1S–BACH1 axis at the rs35976034 locus. We prioritised rs35976034 and implicated *MAP1S* as a context-specific effector in Human coronary artery VSMCs. This is consistent with a lentivirus-based massively parallel reporter assay study in aortic VSMCs that nominated this site as an enhancer for *MAP1S* and supported the link by CRISPRi-mediated repression with reduced *MAP1S* expression.^60^ In our *ex-vivo* system, the rs35976034 CRE gained accessibility with ox-LDL exposure, was ABC-linked to *MAP1S*, and *MAP1S* expression increased concordantly. AlphaGenome further predicted that the alternate ‘T’ allele increases regulatory activity and elevates *MAP1S* expression, consistent with the observed data. Mechanistically, variant-to-motif mapping implicated BACH1 as a putative transcription factor acting at this locus. The alternate ‘T’ allele created a BACH1 motif and showed biased ox-LDL-associated motif usage. Ox-LDL increased *BACH1* and *MAP1S* expression. CRISPR knockout of *BACH1* rescued ox-LDL-induced growth arrest and senescence phenotypes in Human coronary artery VSMCs. BACH1 has been linked to ferroptosis programmes in cardiomyocytes^61^ and vascular endothelial cells^62^, endothelial senescence^63^, and impaired revascularisation after ischaemic injury^64^. However, its role in VSMC dysfunction in atherosclerosis is less clear. Our data therefore support BACH1 as a lipid-stress regulator in coronary VSMCs and nominate a MAP1S-BACH1 circuit as a plausible mediator of rs35976034-linked regulatory effects.

### Limitations

This study provides a multi-omics framework to prioritise ox-LDL-responsive regulatory variants and nominate candidate effector genes at CAD loci. However, some limitations should be noted. First, our human mechanistic work was necessarily performed in a primary human coronary VSMC cells which may not capture the cellular diversity, haemodynamic cues, and immune–stromal interactions of plaques and cannot model the evolution of a disease that occurs over decades. To address this, we tested whether ox-LDL-linked transcriptional programmes correlated with those in human diseased tissue. We meta-analysed independent coronary plaque versus non-plaque single-cell transcriptomes. Ox-LDL-associated signatures were concordantly enriched *in vivo*, including in pre-foam VSMC subpopulations, supporting the physiological relevance of the pathways we nominate. Second, our integrative prioritisation identifies candidate causal variants, enhancers, and effector genes, but therapeutic translatability remains uncertain. Future work could validate variant–enhancer–gene mechanisms and interventions *in vivo*, define their contribution to lesion progression and instability, and test whether modulating these pathways yields therapeutic benefit.

## CONCLUSIONS

In summary, we generated a genome-wide enhancer–gene map of the ox-LDL response in human coronary artery VSMCs by integrating ATAC-seq, H3K27ac, and Hi-C. Ox-LDL induced programmes included lipid handling, autophagy, hypoxia, senescence, and inflammation, while ox-LDL repressed programmes including extracellular matrix and cell-cycle programmes. Using the ABC framework, we linked dynamic distal CREs to 2,008 differentially expressed genes through 4,243 peak-gene connections. The directionality of changes in enhancer accessibility and gene transcription were consistent, supporting enhancer-mediated regulation of the ox-LDL response. ABC-linked, ox-LDL-responsive enhancers were enriched for CAD variants relative to non-linked enhancers and relative to variants associated with non-vascular inflammatory traits, consistent with coronary artery smooth muscle cell–specific genetic disease mechanisms. Locus-level analyses nominated rs76182758 as a candidate regulatory variant acting through an ox-LDL-responsive enhancer-*GUCD1* mechanism and highlighted a rs35976034-linked *MAP1S*–*BACH1* circuit as a plausible mediator of ox-LDL responses. Together, these maps refine effector-gene assignment at CAD loci and prioritise regulatory and potentially therapeutic targets in coronary artery VSMCs.

## Acknowledgments

The authors thank all healthy volunteers who agreed to participate in this study.

