## Supplemental Figures for "CAD variants act through ox-LDL-induced enhancer remodelling to alter VSMC gene programmes"

### SUPPLEMENTARY FIGURES

#### Concentration titration of ox-LDL

To determine the concentration of ox-LDL for treatment, we performed initial exposure of human coronary artery VSMCs to buffer and ox-LDL at: 10, 20, 50, and 100 µg concentrations. The 100 µg/ml concentration showed the most shift in fluorescence (Figure S1) and was subsequently used.

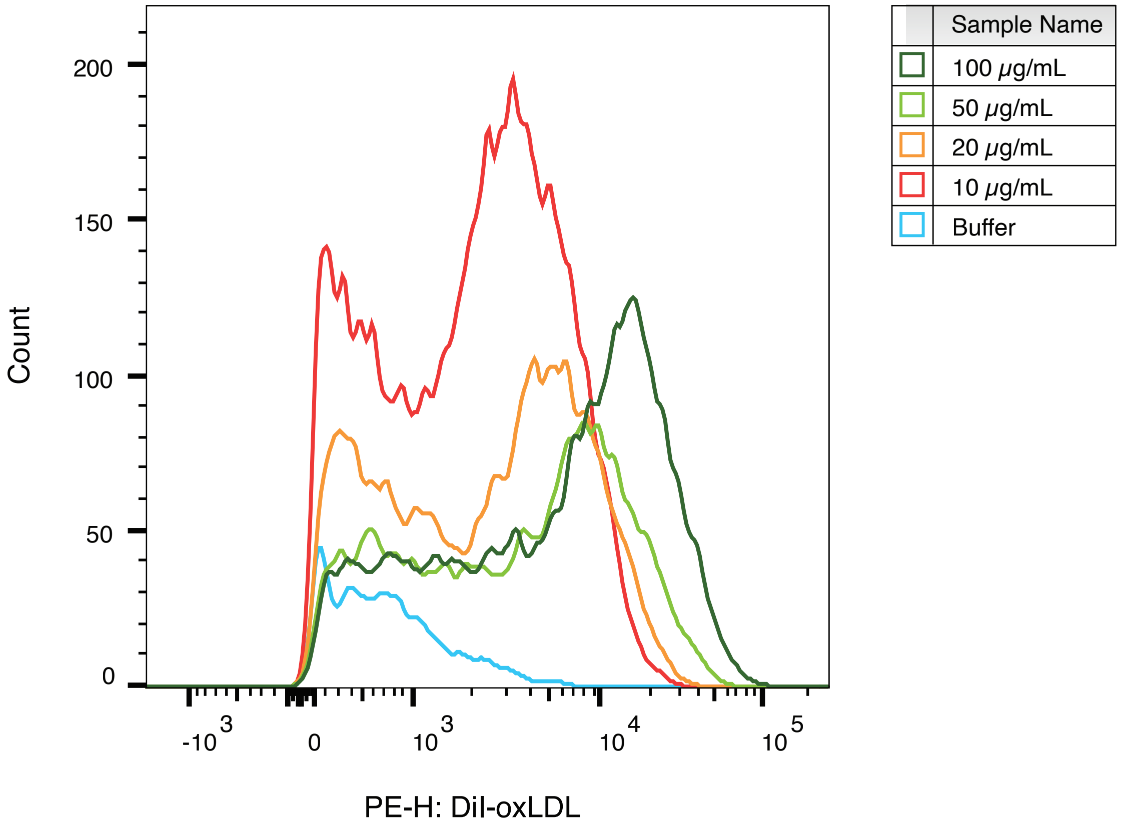

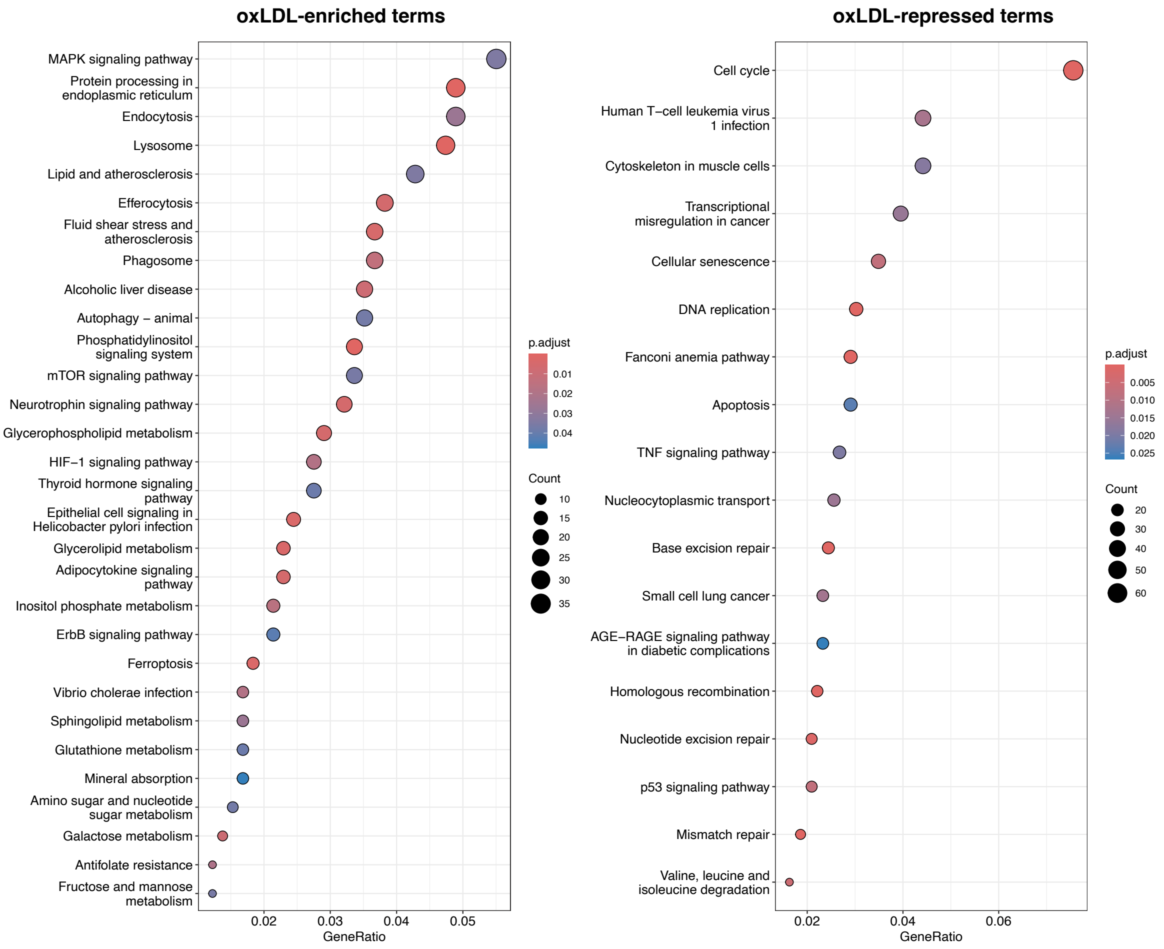


***Fig-S2. Top Kyoto Encyclopaedia of Genes and Genomes (KEGG) terms of ox-LDL transcriptomic signatures***

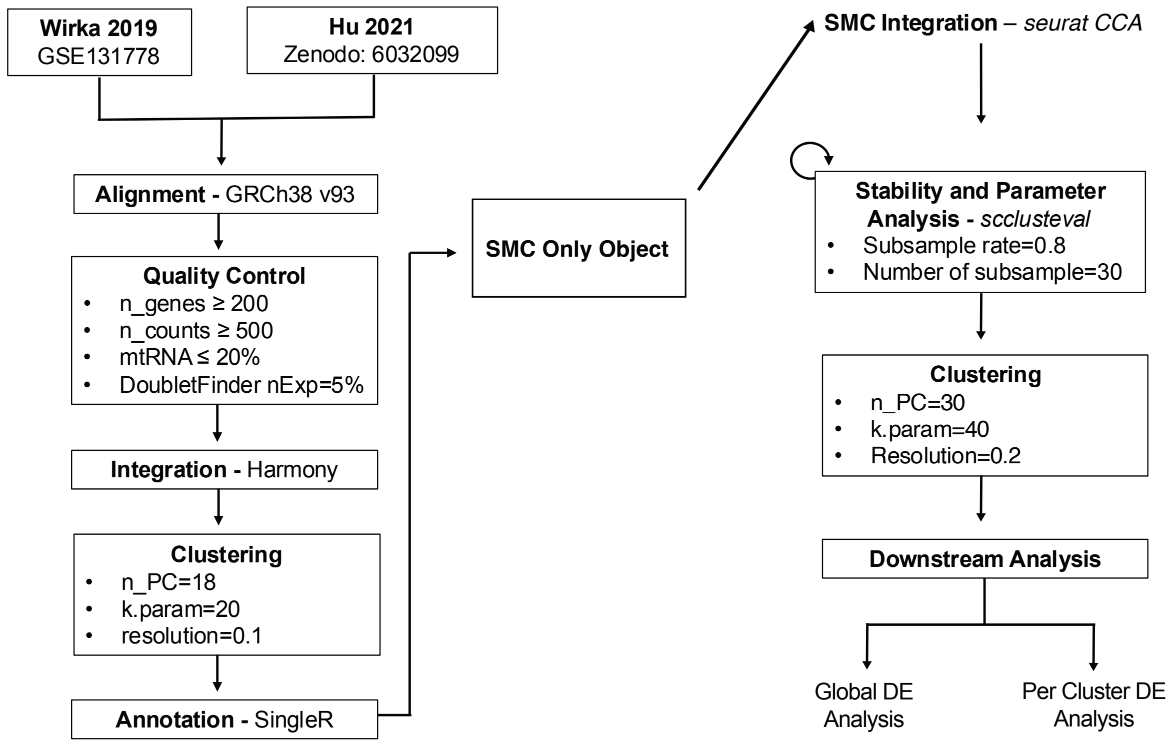


***Fig-S3. Schematic illustration of the methods used for scRNA-seq meta-analysis.***
*See methods for details.*

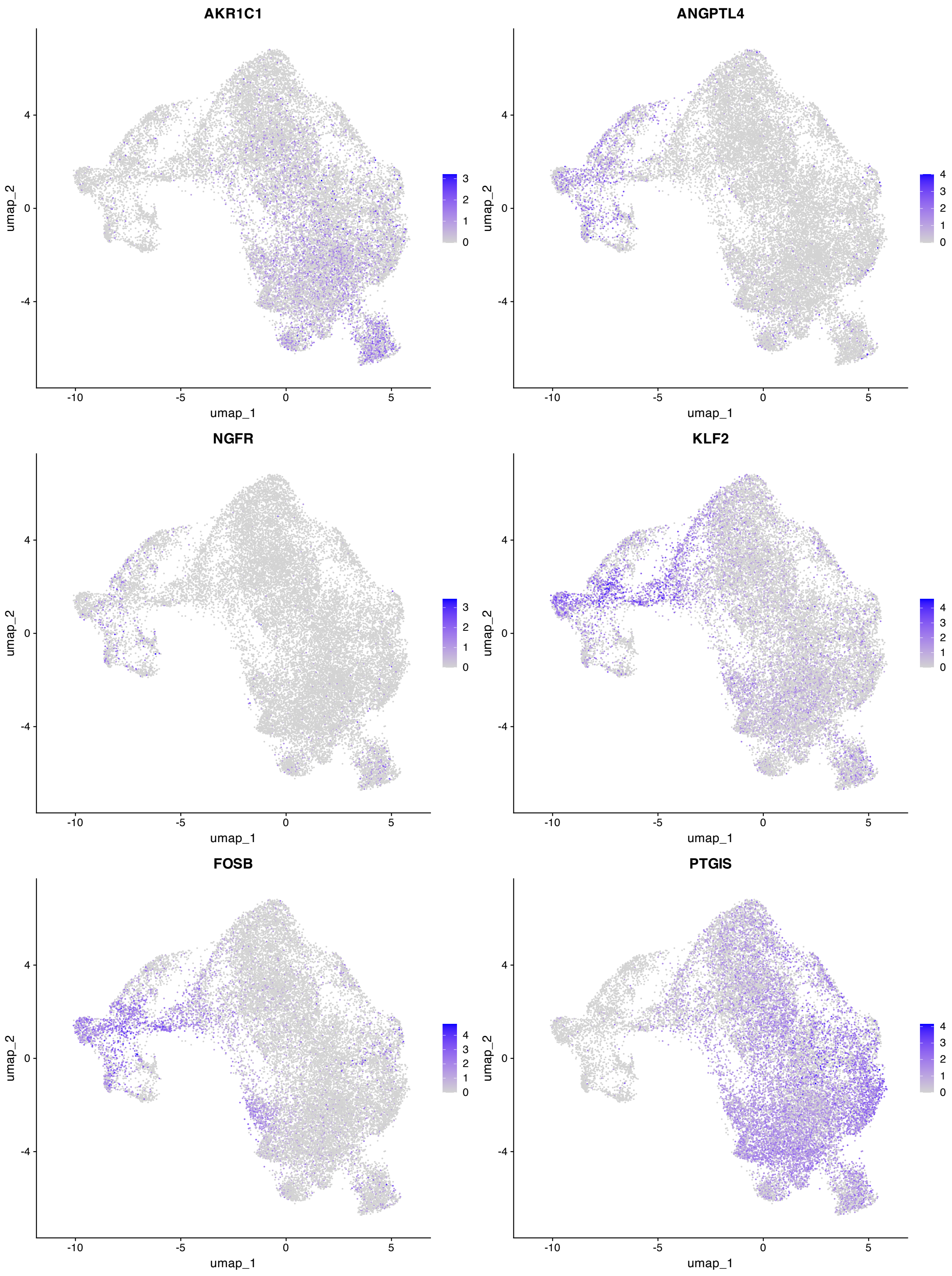


***Fig-S4. Expression of ox-LDL vs Buffer transcriptomic signatures in different VSMC subpopulations in vivo***

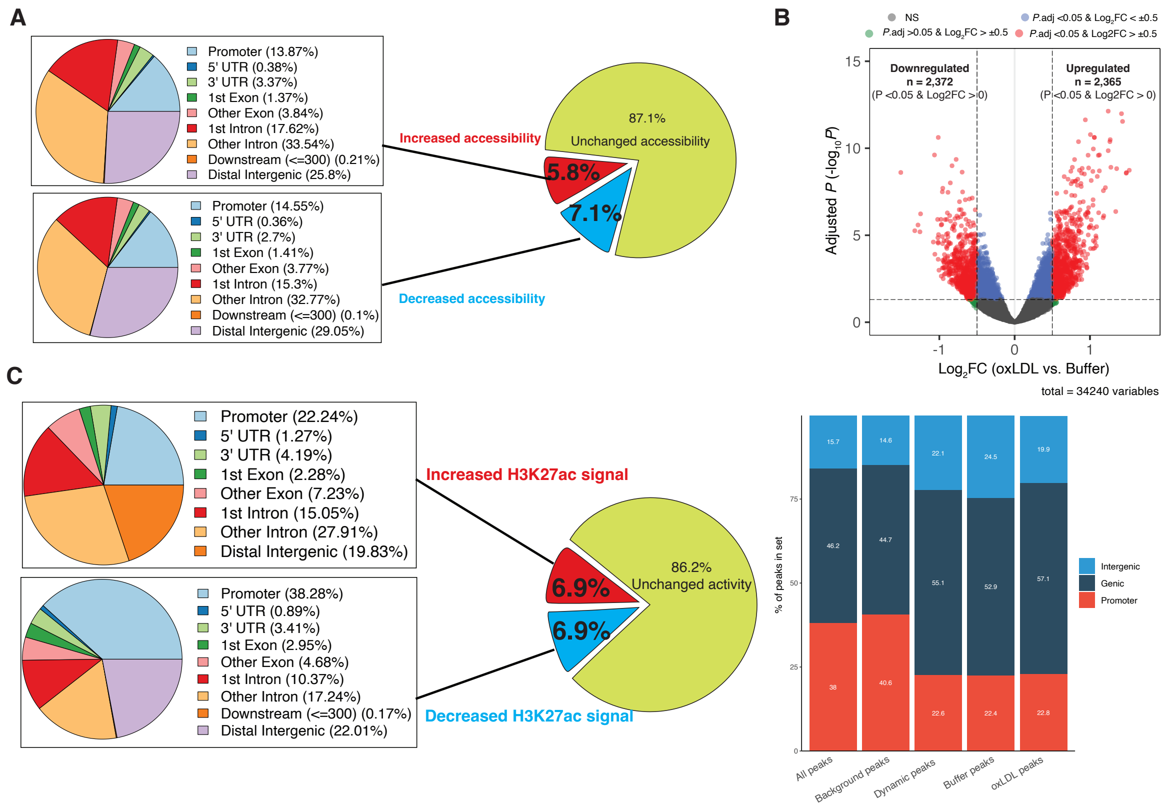


***Fig-S5. Ox-LDL results in significant genome-wide differential accessibility and H3K27ac signals***

***A** Pie charts illustrating the distributions of peaks with increased and decreased accessibility comparing ox-LDL vs. buffer. UTR, untranslated region. **B**, Volcano plot of the H3K27ac signal changes induced by ox-LDL compared to buffer. H3K27ac signal was assessed using chromatin immunoprecipitation with tagmentation sequencing (ChIPmentation). The horizontal dashed line represents P.adj <0.05 and vertical dashed lines are $log_2$ fold change ($log_2FC$) = -0.5 or +0.5. NS, not significant; adjusted P indicates P-values adjusted for multiple comparison using the Benjamini-Hochberg false-discovery rate (FDR) method. **C**, Pie charts illustrating the distributions of peaks with increased and decreased H3K27ac signals by comparing ox-LDL vs. buffer (Left) and proportion of peaks mapping to promoter, genic, and intergenic regions.*

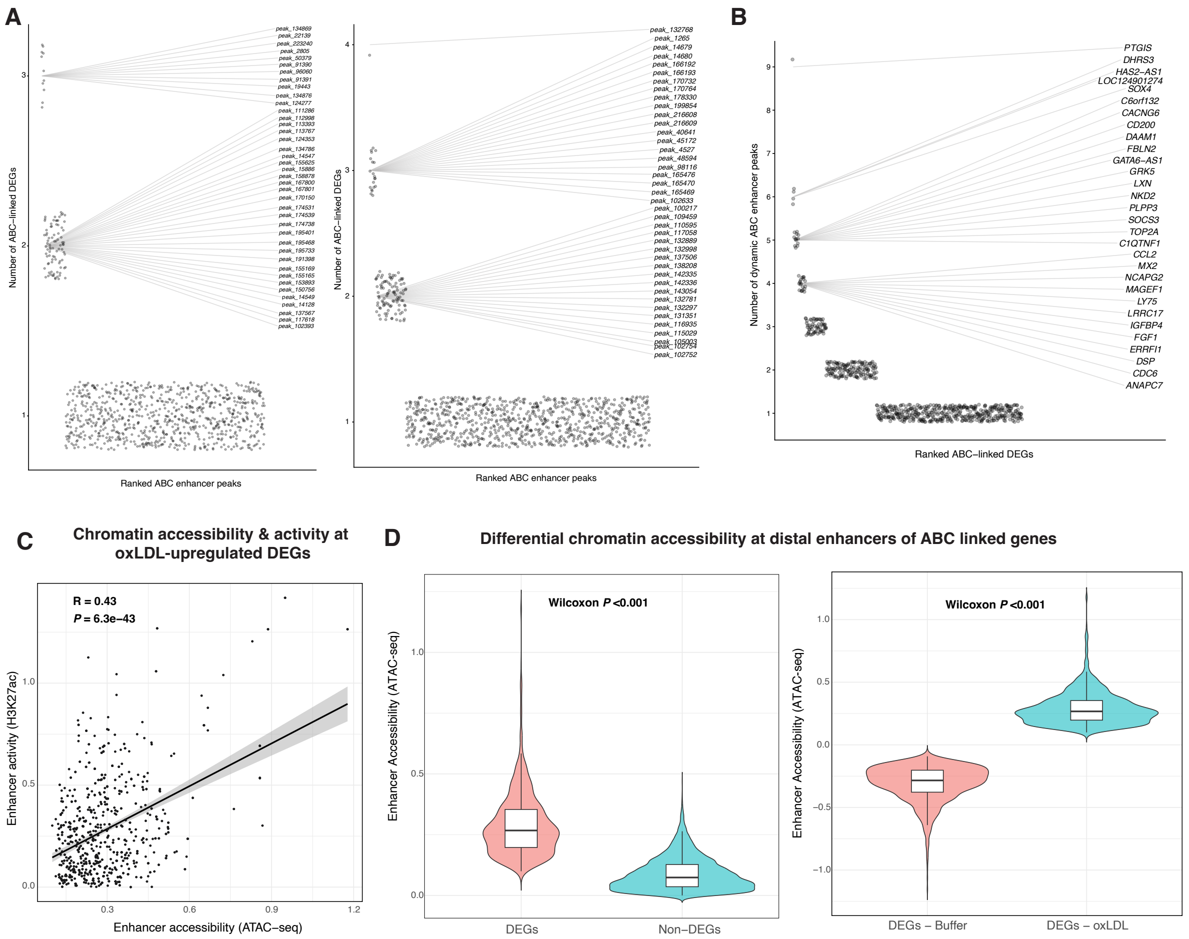


**Fig-S6. ABC enhancer–gene mapping reveals enhancer pleiotropy and gene enhancer burden in ox-LDL-treated Human coronary artery VSMCs. (A)** Enhancer-gene pleiotropy. **Left:** ox-LDL-upregulated dynamic distal enhancers ranked by the number of ABC-linked upregulated DEGs they connect to. **Right:** ox-LDL-downregulated dynamic distal enhancers ranked by the number of ABC-linked downregulated DEGs. x-axis: enhancers (ranked by DEG count). y-axis: number of DEGs per enhancer. **(B)** Ranking of gene-centric enhancer burden. Upregulated ABC-linked DEGs ranked by the number of dynamic distal enhancers linked to each gene. x-axis: genes (ranked by enhancer count). y-axis: number of enhancers per gene. **(C)** Chromatin concordance at dynamic links. Scatter plot of ATAC-seq accessibility vs H3K27ac signal for ox-LDL-up dynamic distal enhancers targeting ox-LDL-up ABC-linked DEGs. x-axis: ATAC-seq signal (or $\log_2$FC). y-axis: H3K27ac signal (or $\log_2$FC). **(D)** Accessibility at enhancers of DE classes. Violin plots comparing enhancer accessibility for distal enhancers linked to ox-LDL-upregulated DEGs versus those linked to non-DEGs and to ox-LDL-downregulated DEGs (statistical test as in Methods; $P < 0.001$).

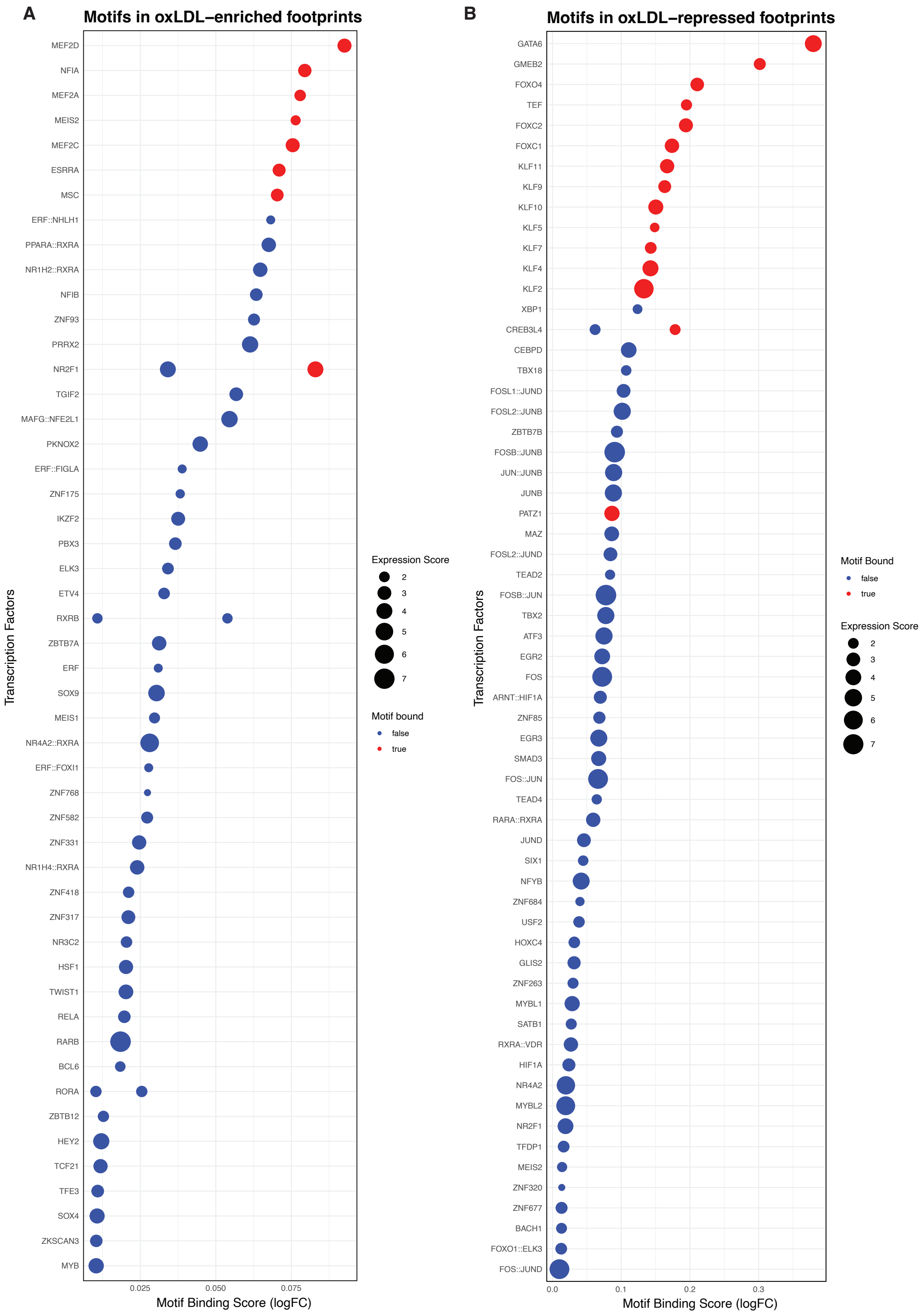


***Fig-S7. Full list of genome-wide transcriptional regulators of the ox-LDL response with significant RNA-seq support. (A)*** *TFs whose motifs show increased footprint occupancy in ox-LDL-enriched footprints (TOBIAS differential binding).* ***(B)*** *TFs whose motifs show increased footprint occupancy in ox-LDL-repressed footprints. All entries*

*pass both criteria: significant differential footprinting (FDR < 0.05; effect thresholds as in Methods) and significant differential expression in RNA-seq (DEG; FDR < 0.05; TMM-normalised). Panels include both concordant and discordant directionality relative to RNA-seq.*

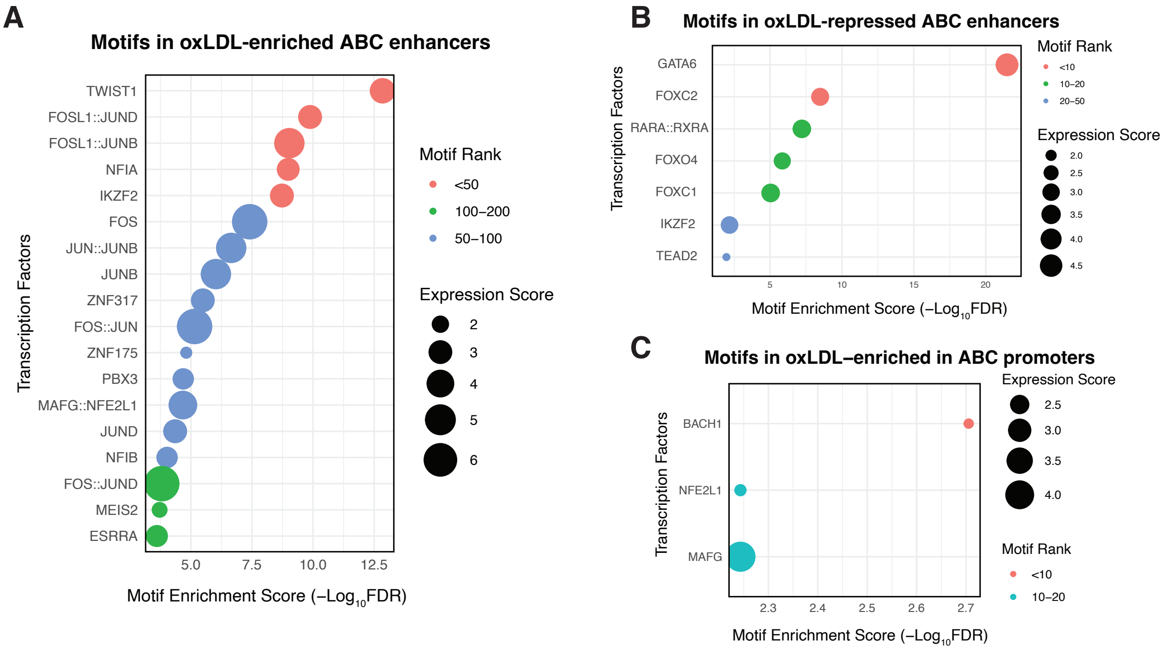


***Fig-S8. ABC enhancer-linked enhancer and promoter motif enrichment with RNA-seq support in Human coronary artery VSMCs. (A)*** *TF motifs enriched in distal enhancer CREs (ABC-linked) whose chromatin become more accessible with ox-LDL versus length/GC-matched background using AME (MEME Suite).* ***(B)*** *TF motifs enriched in distal enhancer CREs (ABC-linked) whose chromatin become less accessible with ox-LDL versus background as in panel (A).* ***(C)*** *Motif enrichment in ABC-linked promoters ('promoters-as-self') overlapping ox-LDL-responsive dynamic CREs, computed with AME; displayed TFs pass the concordance filter (RNA-seq DEG direction matches CRE direction). For panels A–B, entries satisfy both: significant motif enrichment (FDR < 0.05; effect thresholds as in Methods) and significant differential expression in RNA-seq (DEG; FDR < 0.05); panels include both concordant and discordant directionality relative to gene expression. Dynamic CREs are defined by ox-LDL-associated changes in chromatin accessibility. ABC parameters (score threshold, window) and AME settings are detailed in Methods.*

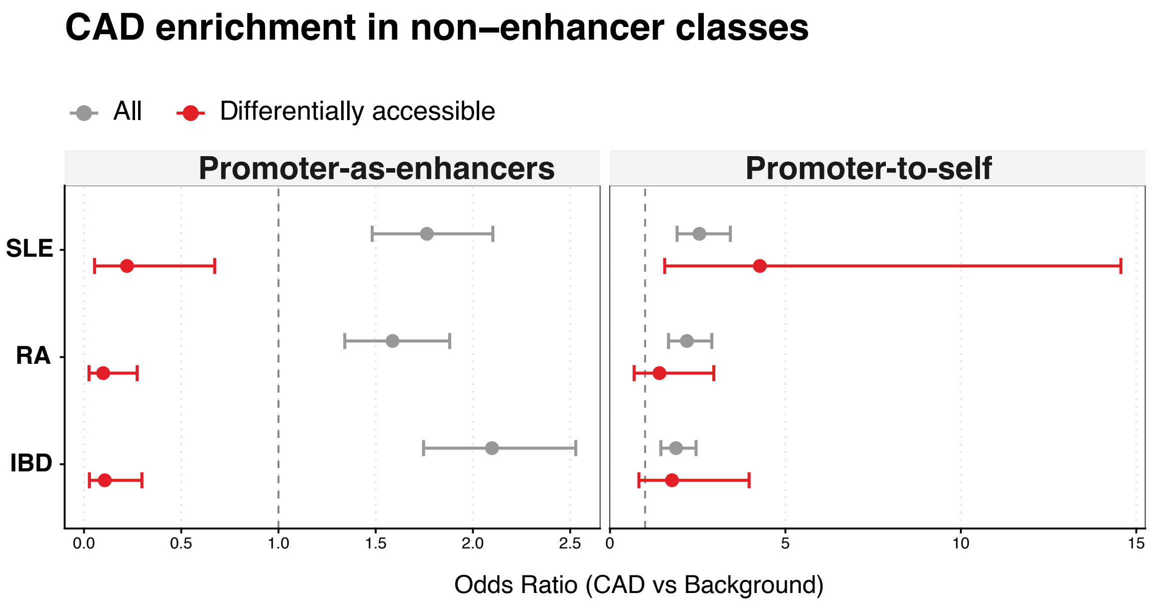


***Fig-S9. CAD variant overlaps in non-enhancer ABC classes.***
*Overlap of CAD variants relative to variants associated with immune disease controls within ABC promoters. Effect sizes are shown for both the full set (All) and the subset of differentially accessible elements. IBD, inflammatory bowel disease; SLE, systemic lupus erythematosus; and RA, rheumatoid arthritis.*

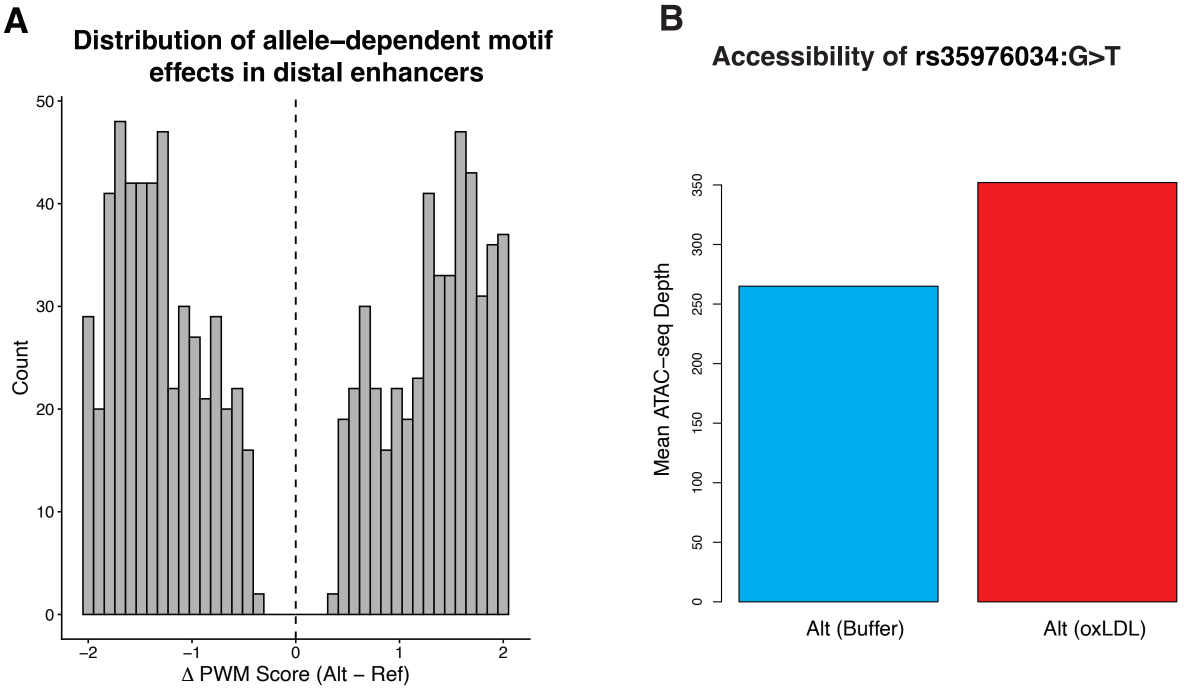


***Fig-S10. Effect of GUCD1 knockout on oxLDL-induced senescence in coronary artery VSMCs.***
***(A)*** *Histogram showing the distribution of ΔPWM values across prioritised variant–motif events, illustrating both positive and negative allele-dependent effects.* ***(B)*** *Descriptive allele-specific accessibility at rs35976034 in a single heterozygous donor. Bar plot*

*shows mean ATAC-seq read depth for the alternate 'T' allele under buffer and oxLDL conditions. Higher alternate-allele depth under oxLDL is consistent with increased accessibility at this locus following oxLDL exposure.*

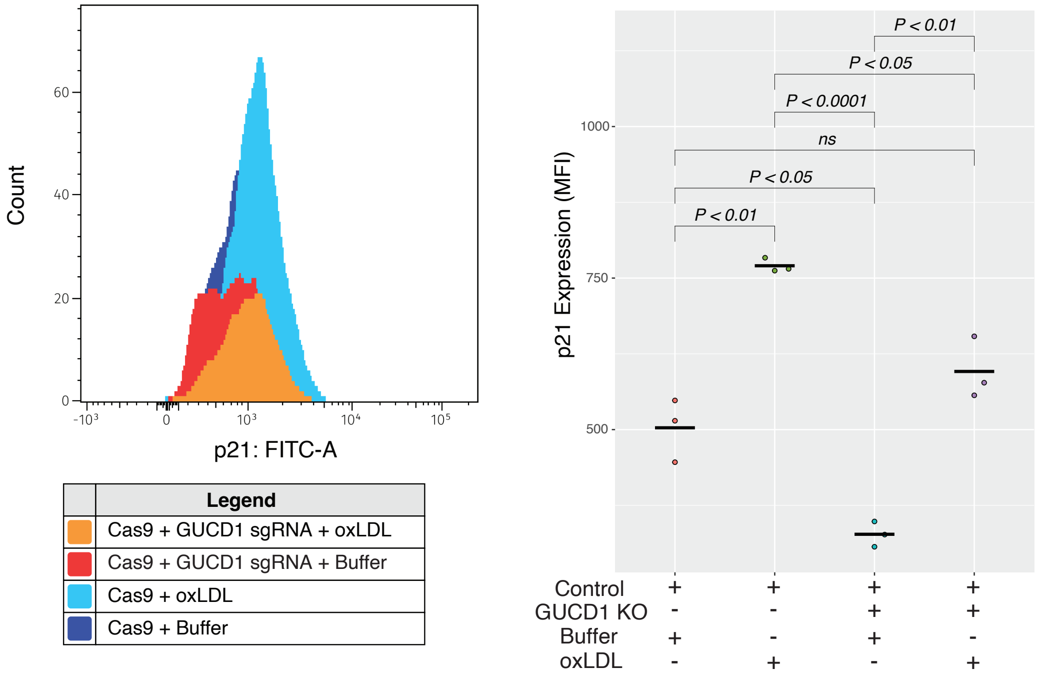


***Fig-S11. Effect of GUCD1 knockout on oxLDL-induced senescence in coronary artery VSMCs.***
*On the left, is the FACS histogram of p21 counts for each condition. On the right, are the quantitative data. See main Figures 6 and 7 for details.*

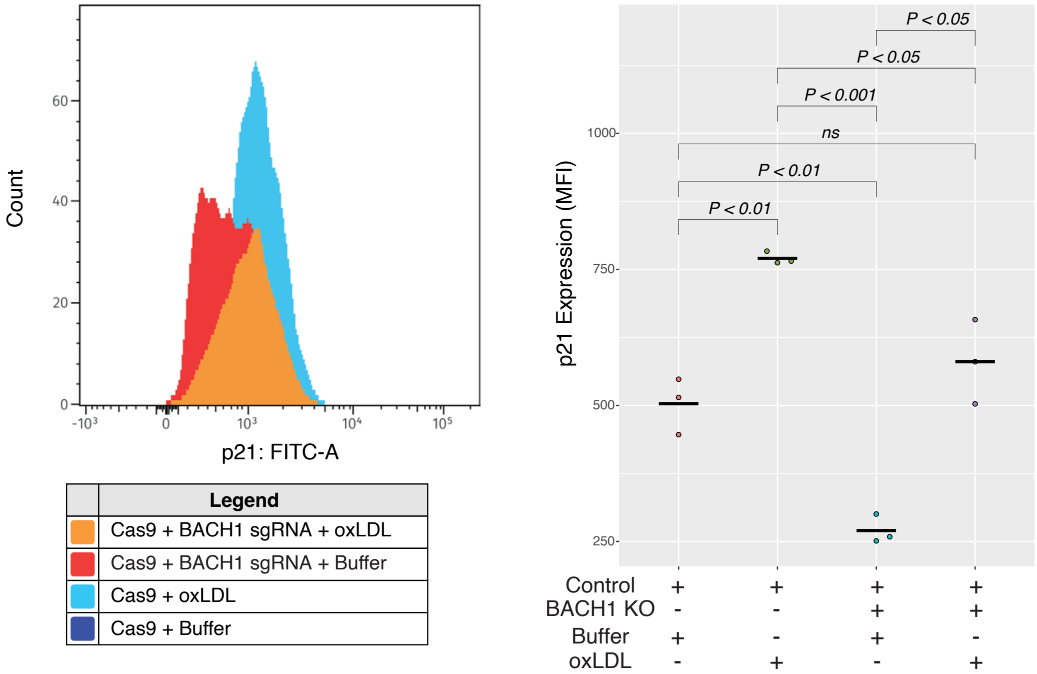


***Fig-S12. Effect of BACH1 knockout on oxLDL-induced senescence in coronary artery VSMCs.***
*Details as in **Fig-S9**.*
